# Optimised dissociation and multimodal profiling of prostate cancer stroma reveal fibromuscular cell heterogeneity with clinical correlates

**DOI:** 10.1101/2025.06.25.661484

**Authors:** Elisabeth Damisch, Elena Brunner, Lukas Nommensen, Lucy Neumann, Georgios Fotakis, Zlatko Trajanoski, Sieghart Sopper, Georg Schäfer, Martin Puhr, Isabel Heidegger, Marianna Kruithof-de Julio, Natalie Sampson

**Affiliations:** Department of Urology, Medical University of Innsbruck, 6020 Innsbruck, Austria; Institute of Bioinformatics, Medical University of Innsbruck, 6020 Innsbruck, Austria; Univ. Clinic for Hematology and Oncology, Internal Medicine V, Medical University of Innsbruck and Tyrolean Cancer Research Institute, 6020 Innsbruck, Austria; Department of Pathology, Neuropathology and Molecular Pathology, Medical University of Innsbruck, 6020 Innsbruck, Austria; Department for BioMedical Research, Urology Research Laboratory, University of Bern, Bern, Switzerland; Department of Urology, Inselspital, Bern University Hospital, University of Bern, Bern, Switzerland

**Keywords:** cancer-associated fibroblast, prostate cancer, single cell RNA sequencing, smooth muscle cell, tissue dissociation, tumour microenvironment

## Abstract

**Background:** Dynamic remodelling of the tumour microenvironment (TME) plays a central role in prostate cancer (PCa) progression, immune evasion and therapy resistance. However, the co-existence of both tumour-promoting and tumour-restraining stromal elements necessitates extensive characterisation of the TME for effective targeting. Fibromuscular cell heterogeneity in PCa remains poorly characterised, in part due to challenges in isolating cells embedded within the desmoplastic stroma. This study therefore aimed to better characterise fibroblast and smooth muscle cell (SMC) populations as the major tissue-resident stromal cell subtypes within the PCa TME.

**Methods:** A PCa single-cell RNA sequencing (scRNA-seq) dataset was re-analysed to define fibromuscular subtypes. Due to low fibroblast yields, an optimised tissue dissociation protocol was developed and benchmarked against two commercial kits via flow cytometry, immunostaining of clinical specimens and *ex vivo* culture. Dimensionality reduction and clustering were applied to the CD31^−^ stromal fraction using a multiparameter surface marker panel. Annotation of the resulting clusters based on their surface marker profile was supported by integrating scRNA-seq and immuno-histological findings.

**Results:** The optimised protocol yielded over twice the viable cells/mg tissue compared to two commercial kits, preserved surface marker integrity, enhanced successful cultivation of mesenchymal cells and recovered diverse stromal subpopulations from benign and malignant samples. Dimensionality reduction and clustering of flow cytometry counts identified 11 distinct CD31^−^ stromal populations. Integration with transcriptomic data and immunofluorescence of clinical specimens identified spatially- and prognostically-distinct fibroblast subtypes, including inflammatory and myofibroblastic cancer-associated fibroblasts, pericytes linked to poor prognosis and a novel SMC subset associated with stromal activation.

**Conclusion:** This study presents a robust workflow for improved isolation and characterisation of fibromuscular stromal cells in PCa. The multimodal approach enabled refined characterisation of phenotypically distinct and clinically-relevant stromal subpopulations within their spatial context providing a foundation for future TME-targeted therapies.

## Introduction

Stromal activation is a dynamic process occurring early in the pathogenesis of prostate cancer (PCa), already evident in pre-neoplastic lesions like prostatic intraepithelial neoplasia (PIN), and plays a critical role during tumour progression, immune evasion and therapy resistance (Pakula et al., 2024; Pederzoli et al., 2023). Thus, the tumour microenvironment (TME) is widely considered a promising therapeutic target. The TME however is highly heterogeneous and contains both tumour-suppressive and -promoting cellular entities, which differentially influence prognosis and therapeutic response (Liu et al., 2023; Pakula et al., 2024).

Prostate interstitial smooth muscle cells (PSMC) are the most abundant cell type in the benign stroma. Beyond their contractile function, PSMC support organ homeostasis and provide a physical barrier that restrains tumour progression (Pederzoli et al., 2023; Thomas et al., 2024; Zhang et al., 2003). In PCa however, PSMC undergo dissociation and secretory activation followed by degeneration and elimination via poorly defined mechanisms (Taboga et al., 2008). Mural cells (vascular SMC (VSMC) and pericytes) in contrast maintain vessel integrity, elasticity and contractility (Muhl et al., 2020). During tumour progression however, cancer cells promote pericyte dissociation resulting in vessel leakage, which facilitates invasion and metastasis (Pederzoli et al., 2023). Moreover, in response to injury or tumour-derived signals pericytes differentiate into myofibroblasts, VSMC or cancer-associated fibroblasts (CAF) (Muhl et al., 2020; Murray et al., 2014; Pederzoli et al., 2023).

Such plasticity is also a hallmark of fibroblasts, which maintain homeostasis in healthy tissues but differentiate into extracellular matrix (ECM)-producing myofibroblasts during wound healing and fibrosis (Muhl et al., 2020; Pederzoli et al., 2023). In PCa, fibroblast abundance increases, whereby their activation modulates the TME via paracrine signalling, ECM remodelling and immune regulation (Liu et al., 2023; Pakula et al., 2024). CAF exhibit a spectrum of interconvertible activation states that are governed by spatial context, biophysical/- chemical properties of the TME, tumour genotype and cellular origin (Jenkins et al., 2022; Pakula et al., 2024). Antigen-presenting, inflammatory and myofibroblastic CAF (apCAP, iCAF and myCAF, respectively) constitute the three major and functionally-distinct CAF phenotypes identified to date (Galbo et al., 2021; Liu et al., 2023). The prognostic implications of CAF states vary by cancer type (Galbo et al., 2021; Jenkins et al., 2022). In PCa, iCAF are abundant in PIN and less aggressive tumours, associated with favourable outcome and thus potentially represent an early/less-activated state, whereas myCAF predominate high grade tumours and are associated with poor prognosis (Brunner et al., 2025; Liu et al., 2023). Given the functional and prognostic implications of CAF heterogeneity and tumour-suppressive action of PSMC, greater characterisation of fibromuscular subpopulations in PCa is critical for the design of therapeutic strategies that specifically target onco-supportive stromal entities and/or maintain tumour-suppressive stromal elements.

Efficient tissue dissociation is essential when studying cellular heterogeneity, even in the spatial era since cell annotation remains reliant on reference single cell RNA-sequencing (scRNA-seq) datasets due to the limited depth and gene coverage of current spatial platforms (Gulati et al., 2025). Fibroblasts are frequently underrepresented in public scRNA-seq datasets (Lendahl et al., 2022) underscoring the challenge in isolating tissue-resident cells, particularly from desmoplastic tissues, such as PCa. Whilst explant outgrowth permits primary fibroblast isolation, this approach may favour migratory subtypes and induce myofibroblastic traits (Waise et al., 2019). Optimising mechano-enzymatic tissue dissociation may reduce this selection bias and permit more representative sampling of *in vivo* stromal diversity (Henry et al., 2018). Moreover, pre-clinical evaluation of drug efficacy increasingly employs three-dimensional organoid models, whereby inclusion of the stromal component is deemed critical for meaningful translation (Richards et al., 2019). Thus, robust tissue dissociation protocols to recover ECM-embedded cell populations are required for diverse applications and unbiased TME characterisation.

We present a tissue dissociation protocol that significantly improves isolation of stromal cell types from small prostate specimens and demonstrate that stromal composition differs between benign and malignant tissues. Using a multimodal approach, we identify and recover multiple fibroblast cell subpopulations from the prostate TME, including a myCAF population associated with poor outcome and an iCAF population linked to favourable prognosis. Further, we characterise prostatic mural and SMC subtypes, identifying a prognostically-relevant pericyte population and dedifferentiated PSMC that were enriched in activated tissues. Data presented herein enhance our understanding of fibromuscular heterogeneity within the prostate TME and may facilitate the future discovery of novel stromal-specific therapeutic targets.

## Material & Methods

### Reagents

All reagents were from Sigma Aldrich (Vienna, Austria) unless otherwise specified.

### Cell lines and culture

The 22Rv1 prostate cancer cell line was obtained from the American Type Culture Collection (ATCC; Rockville, MD), STR validated and maintained according to the distributor’s instructions. Human primary prostatic fibroblasts were established from patients undergoing radical prostatectomy at the University Hospital of Innsbruck using an outgrowth method as previously described (Sampson et al., 2018). All cell lines and primary cells were cultured in a humidified atmosphere at 37°C with 5% CO2 and routinely tested for mycoplasma contamination.

### Tissue harvesting

Prostate tissue samples were collected from consenting treatment-naïve patients undergoing radical prostatectomy due to organ-confined PCa at the University Hospital of Innsbruck (see Declarations). Patient data is provided in Supp.Table 1. A uropathologist (G.S.) obtained ⌀4 mm biopsy cores from freshly excised resections within 1 h of surgery. A small tissue section from one end of each biopsy core and the surrounding biopsy punch site was formalin-fixed and paraffin-embedded (FFPE) for histopathological evaluation using haematoxylin and eosin (HE) staining and TP63/AMACR dual immunohistochemistry. The remaining biopsy core was transported to cell culture facilities in serum-free Dulbecco’s modified Eagle medium (DMEM) with 1.0 g/L glucose, L-glutamine, sodium pyruvate, 3.7 g/L sodium bicarbonate (PAN-Biotech GmbH, Aidenbach, Germany) and supplemented with 1% penicillin/streptomycin (P/S, 10.000 U/ml Penicillin, 10 mg/ml Streptomycin, PAN-Biotech GmbH).

**Table 1.** Overview of proposed annotation of FC clusters.

| gated cluster | proposed annotation |  | selective markers |  |  |
| --- | --- | --- | --- | --- | --- |
|  |  |  | PDPN | MCAM | additional |
| 6 | SMC/<br>mural cells | VSMC2, PSMC, pericytes | - | + | CD140b<sup>+</sup>/CD90<sup>+</sup> |
| 9 |  | PSMC in fibrotic stromal tissue | + | + | (CD140b<sup>+</sup>)/CD90<sup>-</sup> |
| 10+15 |  | VSMC1 | - | + | CD140b<sup>+</sup>/CD90<sup>-</sup> |
| 11 | fibroblasts | non-activated | - | - | CD140b<sup>+</sup>/CD90<sup>+</sup> |
| 1 |  | early-activated | + | - | FAP<sup>+</sup>/CD140b<sup>+</sup>/CD90<sup>-</sup>/CD105<sup>-</sup> |
| 2 |  | myCAF | + | - | FAP<sup>+</sup>/CD140b<sup>-</sup>/CD90<sup>-</sup>/CD105<sup>+</sup> |
| 8 |  | iCAF | + | - | FAP<sup>+</sup>/CD140b<sup>+</sup>/CD90<sup>+</sup>/CD105<sup>-</sup> |

### Tissue pre-processing

Tissue cores were rinsed in 10 ml Hank’s Balanced Salt Solution (HBSS, Lonza Group AG, Basel, Switzerland), weighed, and minced in a small volume of the appropriate enzyme cocktail in a glass petri dish using two scalpels. The minced tissue was transferred to a 50 ml reaction tube containing the remaining enzyme cocktail and incubated in a waterbath at 37°C and further processed as described below.

### Tissue dissociation with commercial reagents

The Miltenyi Biotec Tumor Dissociation Kit human (Miltenyi Biotec KG, Bergisch Gladbach, Germany) and BD Horizon Dri Tumor & Tissue Dissociation Reagent (BD Bioscience, Vienna, Austria) kits were employed according to the manufacturer’s instructions (version 23-22196 from 12/2020 for BD product number 661563 and product sheet version until lot 5250302868 for Miltenyi product number 130-095-929). Briefly, enzyme cocktails were prepared in DMEM (PAN-Biotech GmbH) as per the manufacturers’ instructions. For the BD Horizon kit, two tissue samples were prepared, each using 5 ml of the enzymatic cocktail provided but incubated either for 30 min (as per the manufacturer’s instructions) or a maximum of 60 min to permit comparison to the other protocols. Minced tissue samples were incubated in the appropriate enzymatic cocktails in a 37°C waterbath and gently agitated every 5-10 min for 30 min (BD 30 min sample) or up to 60 min (all other samples). For the latter, digestion was terminated either when large tissue pieces were no longer visible in any sample or at 60 min, whichever was reached first. Cell suspensions were subsequently strained through a 70 µm cell strainer (Corning®, Szabo-Scandic HandelsgmbH, Vienna, Austria). Remaining tissue fragments were pushed through the strainer with the plunger of a 2 ml syringe (BD Discardit™ II) and strainers rinsed with 5 ml cold Dulbecco’s Phosphate-Buffered Saline (DPBS, GIBCO™, Thermo Fisher Scientific, Vienna, Austria). Samples were centrifuged at 300 x g for 8 min at 4°C. Cell pellets were resuspended in 100 µl flow cytometry buffer comprising DPBS (GIBCO™) with 0.5% BSA and 2 mM EDTA (Carl Roth GmbH, Karlsruhe, Germany), hereafter termed FC buffer.

### Tissue dissociation with the optimised protocol

Minced tissue samples were incubated in 3 ml enzyme cocktail prepared as outlined in Supp.Table 2 in a 37°C waterbath with gentle agitation every 5-10 minutes. Digestion was monitored and terminated when large tissue pieces were no longer visible or at 60 min, whichever was reached first. Cell suspensions were filtered through a 100 µm cell strainer (Corning® Szabo-Scandic HandelsgmbH) into a 50 ml reaction tube. Remaining tissue fragments were pushed through the strainer with the plunger of a 2 ml syringe (BD Discardit™ II) and strainers rinsed with 5 ml cold DPBS (GIBCO™). Samples were centrifuged at 120 x g for 10 min at 4°C. Cell pellets were resuspended in 1 ml 1x TrypLE™ Express Enzyme (GIBCO™) and incubated for 2 min at 37°C. The TrypLE reaction was stopped by adding 3 ml FC buffer. The cell suspension was filtered through a 40 µm cell strainer (Corning® Szabo-Scandic HandelsgmbH), the strainer rinsed with 4 ml FC buffer and cells collected by centrifugation at 350 x g for 10 min at 4°C. Cell pellets were resuspended in 100 µl FC buffer.

### Cell viability assay

5 µl freshly dissociated cell suspension was mixed with an equal volume 0.4% trypan blue solution and loaded onto a Neubauer improved chamber (Assistent®, Unilab Technologies GmbH, Innsbruck, Austria). Viable cells, which excluded the trypan blue dye, were counted with a 10x objective under an Olympus CK2 inverted phase contrast microscope (Olympus Europa).

### Cell diameter analysis

Cell viability assay images were captured using a 10x objective on an Olympus CK2 inverted phase contrast microscope (Olympus Europa, Hamburg, Germany) equipped with a JENOPTIK GRYPHAX® ProgRes microscope camera (JENOPTIK Optical Systems GmbH, Jena, Germany). Image acquisition was performed using JENOPTIK GRYPHAX® software (version 2.0.0.68, JENOPTIK Optical Systems GmbH). The images were subsequently analysed using ImageJ (v1.54g). The scale for measurement was calibrated according to the scale bars present in each image. The experimenter observed three distinct groups of cells, categorised as small, medium, and large. Representative images were used to measure the diameter of cells in each group using the measurement tool in ImageJ. The cells were classified into three categories: small (≤5 µm), medium (>5 ≤10 µm), and large (>10 µm). Based on this classification, four replicates from each protocol were analysed, with cells stratified according to the defined size categories.

### Seeding and cultivation of dissociated tissue cells

10 µl freshly dissociated cell suspension was seeded into a 24-well plate (Costar®, Szabo-Scandic HandelsgmbH) in fibroblast outgrowth medium comprising DMEM containing 1 g/L glucose (PAN-Biotech GmbH) supplemented with 20% foetal bovine serum (FBS Supreme, PAN-Biotech GmbH), 1% P/S (PAN-Biotech GmbH), 1% 1 M HEPES (pH 7.2), 1% antibiotic-antimycotic (GIBCO™) and 1% ciprofloxacin. After two weeks undisturbed cultivation, the medium was changed to fibroblast culture medium comprising DMEM containing 1 g/L glucose (PAN-Biotech GmbH) supplemented with 10% FBS (PAN-Biotech GmbH) and 1% P/S (PAN-Biotech GmbH) and the medium changed every 7 days. Cell growth was monitored each week with a 4x objective under an Olympus CK2 inverted phase contrast microscope (Olympus Europa) equipped with a JENOPTIK GRYPHAX® ProgRes microscope camera (JENOPTIK Optical Systems GmbH). JENOPTIK GRYPHAX® software (version 2.0.0.68, (JENOPTIK Optical Systems GmbH) was employed for image acquisition.

### Flow cytometry

1 µl of Intratect 50 g/l infusion solution (Biotest AG, Dreieich, Germany) was added to the remaining 85 µl freshly dissociated cell suspension. The mixture was briefly vortexed and incubated on ice for 5 min. BD Horizon™ Brilliant Stain Buffer Plus (BD Bioscience) and the selected antibodies were added at the indicated concentrations (Supp.Table 3) and samples incubated for 30 min at 4°C protected from light. Samples were made up to a final volume of 2 ml with BD Pharm Lyse™ Lysing Buffer (BD Bioscience) pre-equilibrated to room temperature. After vortexing and incubation for 10 min at room temperature, cells were pelleted by centrifugation at 300 x g for 5 min at 10°C. After washing twice with 2 ml FC buffer, the final cell pellet was resuspended in 100 µl FC buffer and maintained at 4°C until flow cytometry within 1h. At least 3 min prior to measurement, 4 µl BD Pharmingen™ 7-AAD viability dye (BD Bioscience) was added to each sample. Data acquisition was performed on a FACSymphony™ A5 cell analyser (BD Biosciences) equipped with laser lines at 355 nm, 407 nm, 561 nm, and 639 nm, controlled by BD FACSDiva (v9.1) software. Antibodies were titrated beforehand to determine the optimal staining concentration in 100µl cell suspension. Unstained and fluorescence-minus-one (FMO) samples were measured to set gates for the gating strategy during data analysis.

### Analysis of flow cytometry (FC) data

FCS files were analysed using FlowJo™ (BD Life Sciences, v10.10.0). Cells were gated based on size and viability stain (FSC-A/7-AAD) to exclude debris and dead cells. After doublet removal (FSC-A/FSC-H), single viable cells were used to identify leukocytes (CD45), epithelial cells (non-basal: CD326^+^PDPN^−^ and basal: CD326^+^PDPN^+^) and endothelial cells (non-lymphatic: CD31^+^PDPN^−^ and lymphatic: CD31^+^PDPN^+^). Cells not expressing any of these markers were classified as CD31^−^ stroma and used for further analysis of stromal subpopulations. Gates for individual stromal markers were set using unstained and FMO samples. To account for differences in tissue input and overall yield, absolute cell counts per mg of tissue were calculated for each population, enabling direct comparison of isolation efficiency across dissociation protocols. Accordingly, FC analyses are reported as counts per mg of tissue or as percentages of total counts, single viable cells, or parent gates. Absolute numbers were not reported as total cell counts varied greatly between samples and protocols. For the eight patient-matched cohort, histopathologically-validated matched benign (BE = 12) and tumour (CA = 8) samples from eight patients were labelled with keywords for identification and annotation. The CD31^−^ stromal counts from all twenty samples were concatenated into a single file retaining the original sample keywords. No downsampling was applied to avoid data loss and preserve rare stromal subpopulations that may not be equally represented across all samples. Dimensionality reduction was performed using the t-distributed stochastic neighbour embedding (tSNE) option in FlowJo™ based on five stromal markers (MCAM, CD140b, CD90, PDPN, FAP). CD140a and CD105 were excluded since they were expressed in less than 5% (median) of the CD31^−^ stroma and their minimal and inconsistent expression introduced noise, fragmenting the data into small, likely artefactual clusters. The learning configuration was set to auto (opt-SNE), with 1000 iterations, a perplexity of 30, and a learning rate set to 1/12 of the count of the concatenated file. The Exact (vantage point tree) KNN algorithm and the FFT Interpolation (FIt-SNE) gradient algorithm were used. Putative subclusters were calculated with XShift based on five stromal markers (MCAM, CD140b, CD90, PDPN, FAP), using 500 as the number of nearest neighbours (K), the Euclidean distance metric, a subsampling limit of 10^5, and an auto Run ID. To maintain consistency and prevent overclustering, CD140a and CD105 were also excluded in this step, following the same rationale as in the tSNE analysis. Identified clusters were viewed using ClusterExplorer and manual gating. Of the eight patients, the patient 2 BE2 punch was not included in the eight patient-matched cohort but instead used as an FMO control.

### Immunohistochemistry and immunofluorescent staining of archived tissue

For immunohistochemistry, 2 µm FFPE tissue sections were stained using a Ventana Benchmark Ultra automated staining device (Roche Diagnostics GmbH, Vienna, Austria) with the antibodies listed in Supp.Table 3. For multiplex immunofluorescent staining, 2 µm FFPE tissue sections were deparaffinized and rehydrated in a graded alcohol series. Antigen retrieval was conducted via indirect boiling in Dako Target Retrieval Solution pH 9 (Agilent Technologies Österreich GmbH, Vienna, Austria) for 10 min. After cooling, sections were blocked in 3% BSA in Tris-buffered saline (TBS) before being incubated overnight at 4°C with primary antibodies diluted in 0.5% BSA in TBS as specified in Supp.Table 3. Following washing, the sections were incubated with fluorescently-conjugated secondary antibodies (Supp.Table 3) for 1 hour at room temperature. Nuclei were counterstained with 2.5 µg/ml Hoechst 33342 (Invitrogen, Thermo Fisher) before mounting in VECTASHIELD® mounting medium for fluorescence (Vector Laboratories, Inc., Burlington, CA).

### Immunocytochemistry

Cells cultured from dissociated tissues were washed with DPBS (GIBCO™) and incubated with 1 mg/ml collagenase in HBSS for 5-10 min at 37°C. The supernatant and detached cells were collected. Remaining adherent cells were detached via subsequent trypsinization for 5 min at 37°C. The reaction was stopped with fibroblast culture medium. The cell suspension was pooled with collagenase-detached cells before centrifugation at 300 x g for 10 min at 10°C. Cells were seeded onto acid-washed coverslips in a 6-well plate in fibroblast culture medium.

At least 24 h post seeding, cells were rinsed twice with DPBS and fixed in 4% paraformaldehyde in DPBS for 10 min at room temperature. After washing, cells were permeabilized in 0.3% Triton® X-100 (SERVA Electrophoresis GmbH, Heidelberg, Germany) in DPBS for 5 min at room temperature, rinsed and blocked in 1% BSA in DPBS supplemented with 5% donkey serum for at least 1 h. Cells were stained overnight at 4°C with primary antibodies diluted in 1% BSA in DPBS with 0.1% Tween® 20 (SERVA) as specified (Supp. Table 3). After washing with DPBS, cells were incubated for 1 h at room temperature protected from light with fluorescently-conjugated secondary antibodies diluted in 1% BSA in DPBS with 0.1% Tween-20 as outlined (Supp. Table 3). After washing, nuclei were counterstained with 5 µg/ml Hoechst 33342 (Invitrogen, Thermo Fisher) for 15 min at room temperature in DPBS, washed and coverslips mounted in VECTASHIELD® mounting medium (Vector Laboratories).

### RNA in situ hybridization

2 µm FFPE prostate tissue sections were stained via duplex RNA in situ hybridization using the probes indicated (Supp. Table 3) with the RNAscope^TM^ 2.5 High Definition Duplex assay kit (Advanced Cell Diagnostics Inc., Newark, CA) according to the manufacturer’s instructions. Positive (PPIB and POLR2A) and negative (dapB) control probes were hybridized in parallel for all experiments.

### Imaging

Images were acquired using a Zeiss Axio Imager Z2 microscope (Zeiss, Vienna, Austria), equipped with a Pixelink PL-B622-CU camera for brightfield imaging and a monochrome pco.edge 4.2LT camera for fluorescence imaging. TissueFAXS® software (version 7.137, TissueGnostics® GmbH, Vienna, Austria) was employed for image acquisition using a 4x, 10x or 20x air objective; or 40x oil or 63x oil objective, maintaining constant image acquisition settings. For fluorescent images, single channel monochrome images were merged and pseudo-coloured for visualisation as described in the corresponding figure legend with ImageJ (v1.54g).

### Bioinformatic re-analysis of scRNA-seq data

Bioinformatic analyses were conducted in R (v4.2.1) using RStudio (v2022.7.1.554) (Posit team, 2022) incorporating both base R functionalities (R Core Team, 2022) and Bioconductor packages (Huber et al., 2015). Data manipulation was primarily performed with the tidyverse package (v2.0.0) (Wickham et al., 2019).

#### Re-analysis of fibromuscular cell types, cluster identification and annotation

To unbiasedly group cells, principal component analysis (PCA) was performed on highly variable genes using graph-based clustering in the FindClusters function of the Seurat package (Satija et al., 2015). Cluster results were visualized using UMAP plots to verify all visually identified clusters were captured and not under-partitioned. Over-partitioned clusters representing the same biological phenotype were merged into a single cluster. Fibroblast and SMC/mural cell clusters were annotated based on human orthologs of canonical fibroblast and SMC/mural cell marker genes (Muhl et al., 2020) as described (Heidegger et al., 2022) and validated using fibroblast and SMC gene signatures from healthy human prostate (Henry et al., 2018). Fibroblast and SMC/mural cell clusters were separately clustered and the data re-scaled accordingly. Subclusters were denoted F1-F5 (fibroblast subclusters) and M1-M5 (SMC/mural cell subclusters) (Supp. Table 4).

#### Subcluster marker identification and pathway analyses

Fibroblast and SMC/mural cell subclusters were separately analysed using the FindAllMarkers function of the Seurat package (Satija et al., 2015) (logfc.threshold set to 0.2) (Supp. Table 5). The top 15 most significantly upregulated genes (adjusted P-value (P.adj) <0.05, avg_log2FC > 0) were used as the subcluster-specific marker genes (Supp. Table 6). Pathway analyses were performed using clusterProfiler (v4.6.2) (Wu et al., 2021) and msigdbr (v7.5.1) (Dolgalev, 2022) (Supp. Table 7-8).

#### Gene plotting

Violin-, dot- and UMAP plots were generated using VlnPlot, DotPlot, FeaturePlot and DimPlot functions from the Seurat package (Satija et al., 2015). Median statistics for Violin plots were added using the stats_summary function of the ggplot2 package (v3.4.2) (Wickham, 2016). Heatmaps were plotted with the pheatmap function of the pheatmap package (v1.0.12) (Kolde, 2019). Colour gradients for dotplots or heatmaps were generated with the brewer.pal function of the RColorBrewer package (v1.1-3) (Neuwirth, 2022).

#### Latent time analysis

Spliced and unspliced counts matrices were constructed using the run10x command from the velocyto (v0.17.17) command line tool (La Manno et al., 2018). The human genome (build GRCh38) was used as reference genome. To decrease the risk of confounding factors in the downstream analysis, expressed repetitive elements were masked using the appropriate (build GRCh38) expressed repeat annotation from the UCSC genome browser (Kent et al., 2002). The SeuratDisk R package (v0.0.0.9019) (Satija et al., 2015) was used to convert between Seurat and anndata formats and the resulting files imported to python to carry out the latent time analysis using scVelo (v0.2.5) (Bergen et al., 2020). RNA velocity was estimated by utilizing the dynamical modelling approach as implemented in the scVelo package. Gene-specific latent timepoints obtained from the dynamical model were then coupled to a universal gene-shared latent time, which represents the cell’s internal clock and is based only on its transcriptional dynamics using the scvelo.tl.latent_time() function. The UMAP plot representing the universal latent time was constructed using the scvelo.pl.scatter() function, while the heatmap plot for gene-specific latent times was constructed using the scvelo.pl.heatmap() function.

#### Assignment of gene signature scores

Gene signature scores were calculated with the ScoreSignatures_UCell function of the UCell package (v2.2.0) (Andreatta and Carmona, 2021) using the check_sig function of the hacksig package (v0.1.2) (Carenzo et al., 2022) to ensure that >75% of the signature genes were present in the query dataset (Supp. Table 9). Calculated gene signature scores were added to the Seurat object using the AddModuleScore_UCell function from the Ucell package (Andreatta and Carmona, 2021)and score distribution within each cluster depicted as a Seurat FeaturePlot or DotPlot (Satija et al., 2015).

## Analysis of TCGA data

RNA-seq and clinical data of TCGA prostate adenocarcinoma (PRAD) (Abeshouse et al., 2015) samples were downloaded from UCSC Xena (https://xenabrowser.net/datapages/). Combined z-scores were calculated for bulk transcriptomic- or scRNA-seq-derived gene signatures as described above and compared between clinical parameters. The R package ggsignif (Ahlmann-Eltze and Patil, 2021) was used to perform t-tests. For visualization as heatmaps, only samples with the sample type “primary tumor” were selected. Disease-free survival (DFS) analyses were performed using GEPIA2 (Tang et al., 2019) using the group cut-offs indicated in the corresponding figure legend. scRNA-seq derived signatures were generated from the top 5 upregulated genes ranked according to adjP value except for cluster M5, which used the top 5 genes ranked according to average log2FC. Gene signatures are provided in the Supplemental Table file in Supp. Table 10).

### Statistical analyses

Statistical analyses were performed using GraphPad Prism (v9.5.0 and v10.1.2, GraphPad Software, LLC). Data in plots are shown as the median with the interquartile range. The number of biological replicates (n) is stated in the corresponding figure legends whereby all experiments were independently repeated at least three times. Adjusted P-values <0.05 were considered statistically significant, whereby statistical significance is denoted n.s., not significant; *, P<0.05; **, P<0.01; ***, P<0.001. Outlier detection was conducted using Grubbs’ test (alpha = 0.2) for single suspected outliers or the ROUT method (Q = 1%) for multiple expected outliers. Normality testing was performed on the largest dataset of optimised samples using the Kolmogorov-Smirnov test and applied to all datasets. For the viability assay, comparisons between two groups were performed using the two-tailed unpaired Welch’s t-test, while comparisons among more than two groups were conducted using Brown-Forsythe and Welch ANOVA tests with Dunnett T3 correction for multiple comparisons. For FC data, all parameters except stromal markers were considered dependent and exclusive to the cell population. These were analysed using a mixed-effects model with the Geisser-Greenhouse correction, applying Sidak’s multiple comparisons test for two-group comparisons or Dunnett’s multiple comparisons test for more than two groups. Being independent and not exclusive to the CD31^−^ stroma population, data values from stromal markers were compared using 2-way ANOVA with either Sidak’s or Dunnett’s multiple comparisons test according to the number of groups compared.

## Results

### The prostate comprises multiple distinct fibroblast subpopulations *in vivo*

To better characterise prostate fibromuscular heterogeneity, we re-analysed fibroblast and SMC/mural cells in our previously published scRNA-seq dataset of treatment-naïve localised PCa and patient-matched benign-adjacent samples, which were previously classified only into broad stromal cell types (Heidegger et al., 2022). Re-clustering yielded five fibroblast (F1–F5) and five SMC/mural cell (M1–M5) clusters, with all subclusters detected in each patient (Fig. 1A, Supp. Fig. 1A-B). All subclusters expressed *VIM* and displayed similarity to published fibroblast or SMC/mural cell signatures (Supp. Fig. 1C-E and G), whereby the F and M superclusters were readily distinguished via differential expression of *PDGFRA* vs. *MCAM* in line with previous reports (Fig. 1B-C, Supp. Fig. 1G) (Muhl et al., 2020).

**Figure 1.**
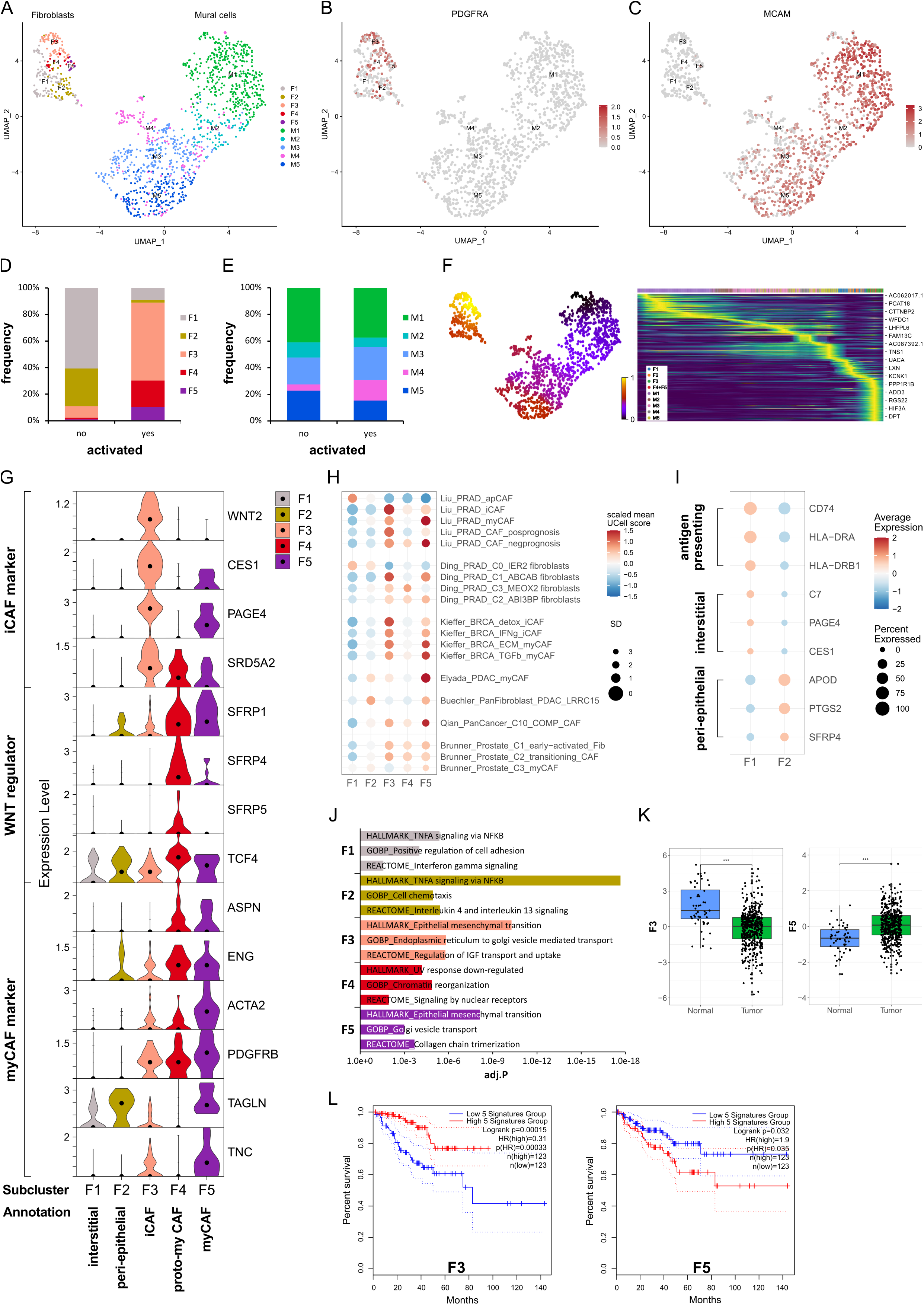
Identification of fibromuscular cell subpopulations in the prostate cancer microenvironment. **(A-J)** Re-analysis of fibromuscular cells in a scRNA-seq dataset (Heidegger et al., 2022) from five patient-matched benign prostate and localised PCa samples. **(A)** UMAP visualisation of fibroblast (F) and SMC/mural cell (M) subpopulations. **(B-C)** Feature plots showing expression of canonical markers for **(B)** fibroblasts (*PDGFRA)* and **(C)** SMC/mural cells (*MCAM)*. **(D-E)** Frequency distribution of **(D)** fibroblast and **(E)** SMC/mural cell subpopulations relative to total fibroblasts and SMC/mural cells, respectively, stratified by histopathological status defined as activated (presence of PIN, inflammation, and/or cancer) or non-activated (absence of these features). **(F)** scVelo latent time modelling of subpopulation fates displayed on the UMAP. **(G)** Violin plot depicting expression levels of selected markers across fibroblast (F) subclusters in the scRNA-seq dataset. **(H)** Expression levels of published scRNA-seq signatures in each fibroblast subpopulation whereby Ucell scores are depicted as scaled mean per subcluster. **(I)** Average expression of top-scoring signature genes for benign human prostate interstitial and peri-epithelial fibroblasts (Joseph et al., 2021) and antigen-presenting markers (Elyada et al., 2019). **(J)** Top upregulated Hallmark, Reactome and Gene Ontology Biological Pathway (GOBP) pathways for each fibroblast subpopulation. **(K)** Expression of F3/F5-specific gene signatures in the TCGA-PRAD cohort as combined z-scores. Statistical significance was determined using the R package ggsignif (Ahlmann-Eltze and Patil, 2021) and is denoted ***, P<0.001. **(L)** Kaplan-Meier curves of DFS in the TCGA-PRAD cohort generated with GEPIA2 using the five most upregulated F3- or F5-specific genes as in (K) and upper/lower quartiles as group cut-off. Dashed lines indicate the 95% CI. Source data for panels (H) and (J-L) are provided in the Source Data file.

Stromal remodelling occurs early during prostate tumourigenesis with “iCAF” subtypes reported in benign-adjacent tissues (Jenkins et al., 2025; Pallares et al., 2006; Tuxhorn et al., 2002). Thus, samples were stratified into “activated” vs. “non-activated” defined as the presence or absence of PIN, inflammation and/or malignant glands in the original tissue sample, respectively. F1 and F2 were most abundant in non-activated samples, displayed low expression of canonical activation markers, such as *PDGFRB* and *ACTA2*, and were placed at the earliest timepoints of the fibroblast trajectory by latent time modelling (Fig. 1D, F-G). Whilst F1 expressed prostate interstitial fibroblast markers (*C7, PAGE4, CES1, RSPO3*), F2 expressed genes associated with prostate peri-epithelial fibroblasts (*APOD, PTGS2, SFRP4*) (Joseph et al., 2021) (Fig. 1I, Supp. Fig. 1F). Duplex *in situ* hybridisation (dISH) confirmed that cells expressing F2 markers *APOD* and *SFRP4* were largely distinct from those expressing the F1 marker *C7* in benign-adjacent tissues (Supp. Fig. 2A) with *APOD^+^*/*SFRP4^+^* cells typically in close proximity to epithelial glands whereas *C7^+^* cells were frequently more distal, particularly within the interstitial stroma (Supp. Fig. 2A). F1 also exhibited features ascribed to apCAF (Fig. 1H-I, Supp. Fig. 1F) and displayed enrichment of pathways associated with antigen-processing (Fig. 1H-J, Supp. Table 8) consistent with a non-professional antigen presenting capacity of fibroblasts under physiological conditions (Harryvan et al., 2021).

F3–F5 were annotated as CAF due to their expression of activation markers (e.g. *PDGFRB, ACTA2*), enrichment in histo-morphologically activated tissues, and positioning at later timepoints in the fibroblast trajectory (Fig. 1D, F-G). F3 and F4 differentially expressed *C7* and *APOD*, suggesting they may represent activated counterparts of interstitial (F1) and peri-epithelial (F2) fibroblasts, respectively (Supp. Fig. 1F). Supportively, in malignant tissues F3 marker *WNT2* was co-expressed with *C7* whereas F4 marker *SFRP4* was co-expressed with *APOD* (Supp. Fig. 2B). Moreover, cells abundantly expressing *SFRP4* expressed little *C7* and primarily displayed a peri-epithelial distribution (Supp. Fig. 2A-B). Conversely, cells expressing abundant *C7* co-expressed little *SFRP4* and were interspersed throughout the stroma. F5 however expressed *C7*, *APOD*, and *SFRP4*, the latter two albeit at lower levels than F4 (Supp. Fig. 1F). Indeed, in addition to cells expressing a preponderance of either *C7* or *APOD* as observed in benign-adjacent tissues (Supp. Fig. 2A), fibroblasts co-expressing *SFRP4* and *C7* or *C7* and *APOD* were observed throughout the tumour-associated stroma (Supp. Fig. 2B), implying that F5 may represent a later activation state common to both peri-epithelial and interstitial fibroblasts, as reported in other solid tumours (Croizer et al., 2024; Hanley et al., 2023).

F3 expressed iCAF-associated genes (*WNT2, CES1, IGF1, SELENOP*) and was enriched for the iCAF-associated pathway coagulation as well pathways related to insulin-like growth factor transport, endoplasmic reticulum transport and epithelial-mesenchymal transition (EMT) (Fig. 1G and J, Supp Fig. 1G, Supp. Table 8). Consistently, F3 exhibited high similarity to several iCAF-associated signatures, including PCa iCAF, detox iCAF and IFNg iCAF from breast cancer (Kieffer et al., 2020; Liu et al., 2023) (Fig. 1H, Supp. Fig. 1H). F3 also expressed the prostate-related markers *PAGE4* and *SRD5A2*, which are involved in androgen receptor (AR) regulation and testosterone metabolism, respectively (Audet-Walsh et al., 2017; Sampson et al., 2012) (Fig. 1G). Annotation of F3 as iCAF was further consistent with our recent study demonstrating expression of CES1 and PAGE4 by C1 prostate fibroblasts exhibiting an iCAF-like phenotype and in PIN and low Gleason tumours (Brunner et al., 2025).

F4 was delineated by upregulation of several WNT regulators, including *SFRP4*, with WNT/beta-catenin signalling among the top enriched GO biological pathways (Fig. 1G, Supp. Table 8). While *SFRP4^+^* CAF subsets have been reported in several cancer types (Andersen et al., 2024; Ding et al., 2024; Du et al., 2024; Kieffer et al., 2020; Ning et al., 2024), F4 showed limited similarity to published iCAF/myCAF signatures (Fig. 1H, Supp. Fig. 1H). However, modest co-expression of myCAF-associated markers *ASPN* and *ENG* implied F4 may denote an intermediate CAF substate concordant with reports that myofibroblast transition proceeds via a WNT-dependent pathway (Cohen et al., 2024) (Fig. 1G). F4 was thus annotated as proto-myCAF.

Of all fibroblast clusters, F5 expressed the highest levels of canonical myCAF markers (*ASPN, ENG, PDGFRB, ACTA2, TAGLN, TNC, ITGA11, CTHRC1*) and was enriched for myCAF-associated pathways, such as collagen chain trimerisation and EMT (Fig. 1G, J). F5 also demonstrated strong similarity to published myCAF signatures, including Elyada myCAF and LRRC15^+^ fibroblasts from pancreatic cancer, pan-cancer C10_COMP^+^ CAF, ECM/TGFβ myCAF from breast cancer and Liu myCAF from PCa (Buechler et al., 2021; Elyada et al., 2019; Kieffer et al., 2020; Liu et al., 2023; Qian et al., 2020) (Fig. 1H, Supp. Fig. 1E and H). We also noted similarity of F5 with late-activated C3/myCAF from PCa, which we recently reported display functional myCAF hallmarks with the defining markers ITGA11 and ENG abundantly expressed in aggressive but not low-grade PCa (Brunner et al., 2025). F5 was thus annotated as myCAF.

The prognostic implications of fibroblast phenotypes vary across cancer types. F3 markers, indicative of the iCAF phenotype, inversely correlated with Gleason score, biochemical relapse, T and N stage and were positively associated with disease-free survival (DFS) (HR 0.31; *P*=0.00015) (Fig. 1K-L, Supp. Fig. 3A-D). Conversely, F5 markers, representative of the myCAF phenotype, were significantly upregulated in tumour samples, positively associated with T stage and negatively associated with DFS (HR 1.9; *P*=0.035) (Fig. 1K-L, Supp. Fig. 3A-D). Collectively, these findings indicate the prostatic stroma harbours multiple fibroblast subpopulations with prostate iCAF and myCAF substates positively and negatively associated with clinical outcome, respectively.

### Stromal activation is associated with changes in smooth muscle cell subpopulations

While the tumour suppressive role of PSMC is well established, prostatic SMC and mural cells remain poorly characterised at the molecular level. Of all M subclusters, M1 and M2 most abundantly expressed contractile markers (*MYL9*, *MYLK, MYH11, TAGLN* and *CNN1*) and were enriched for myogenesis and smooth muscle contraction gene sets (Fig. 2A-C, Supp. Fig. 1G and I). M1 expressed the highest level of these markers and was further delineated by expression of *KCNAB1* and *RERGL*, reportedly specific to arterial VSMC (Barnett et al., 2024) (Fig. 2A, Supp. Fig. 1G and I). Indeed, CNN1 expression was highest in the outermost VSMC layer of arteries and medium-sized (50-400 µm diameter) vessels consistent with distinct VSMC phenotypes residing within the vascular media (Frid et al., 1997) (Supp. Fig. 1J-K). In addition, M1 and M2 abundantly expressed the Ca^2+^ regulator *PLN,* which localised to the multi-layered MCAM^+^ walls of CD31^+^ vessels (Fig. 2E and Supp. Fig. 1K) leading us to annotate M1 and M2 as two distinct VSMC subtypes (VSMC1 and VSMC2).

**Figure 2.**
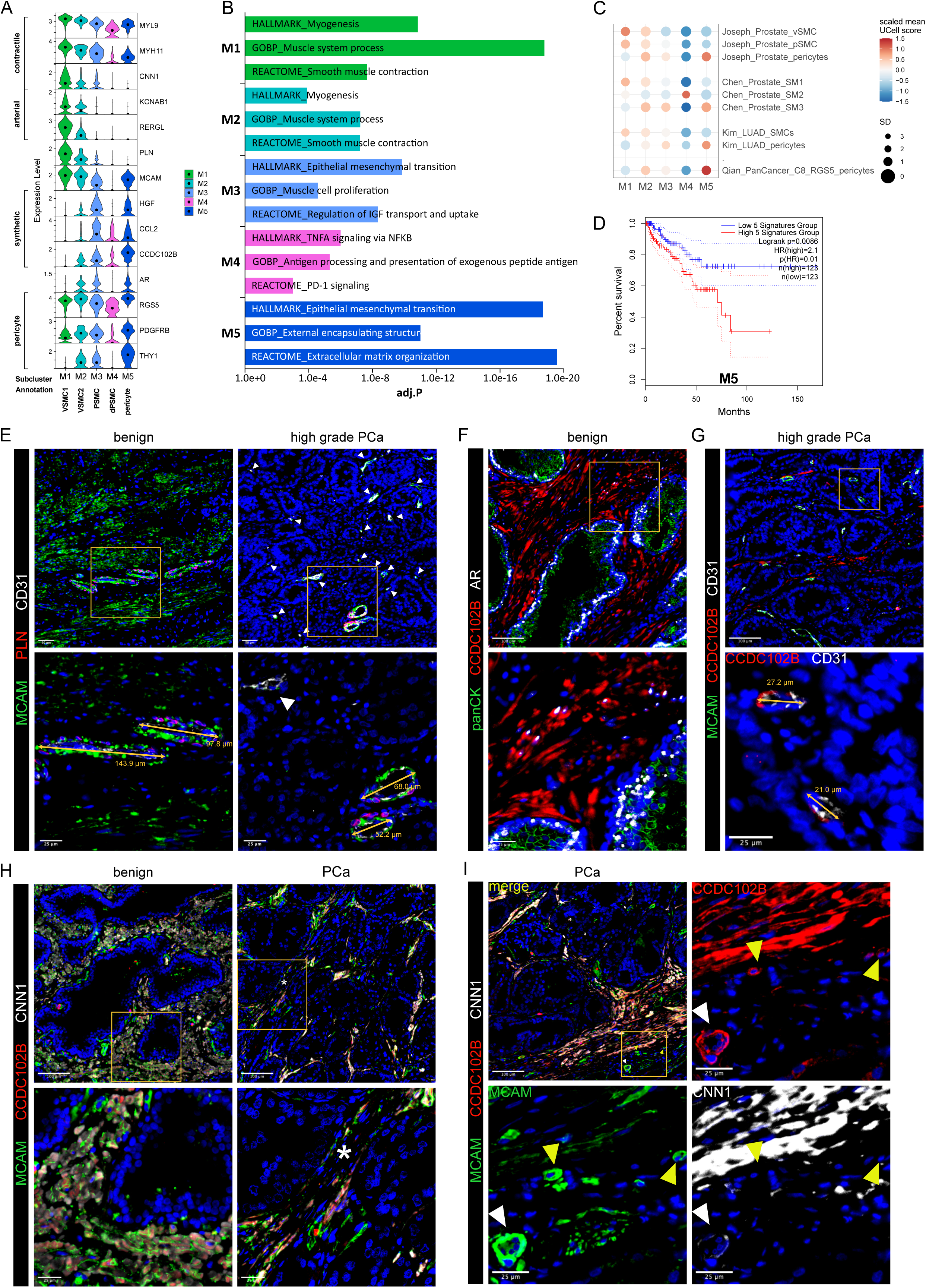
Differential expression of PLN, CCDC102B and CNN1 distinguish prostate PSMC and mural cell subtypes. **(A-D)** Re-analysis of SMC/mural cell subclusters in a scRNA-seq dataset (Heidegger et al., 2022) from five patient-matched benign prostate and localised PCa samples. **(A)** Violin plot depicting expression levels of selected genes demarcating distinct subclusters, which were subsequently annotated as vascular SMC (VSMC), prostate interstitial SMC (PSMC), dedifferentiated PSMC (dPSMC) and pericytes. **(B)** Top upregulated Hallmark, Reactome and Gene Ontology Biological Pathway (GOBP) pathways for each subpopulation. **(C)** Expression of published scRNA-seq signatures across M subclusters whereby Ucell scores are depicted as scaled mean per subcluster. **(D)** Kaplan-Meier curves of DFS in the TCGA-PRAD cohort generated with GEPIA2 using the five most upregulated M5-specific genes and upper/lower quartiles as group cut-offs. Dashed lines indicate the 95% CI. **(E-I)** Immunofluorescent staining of human prostate tissue sections of indicated pathology and patient-matched benign-adjacent areas using the antibodies indicated whereby font colour denotes pseudo-colouring in the displayed merged images. Nuclei were counterstained using Hoechst 33342 (blue). **(E-H)** Enlarged images of orange boxed regions are shown beneath the parental image. **(E)** Cells positive for the M1/M2-enriched marker PLN co-express MCAM and surround CD31^+^ vessels in benign-adjacent tissue (*left)* whereas MCAM^+^ PSMC lack PLN. Large but not smaller CD31^+^ blood vessels (white arrowheads) in high-grade PCa *(right)* are similarly surrounded by a layer of PLN^+^MCAM^+^ VSMC. Decreased MCAM immunopositivity is observed in PSMC but not VSMC in high-grade PCa. Due to strong differences in MCAM expression levels between VSMC and PSMC, enlarged regions were acquired using shorter exposure times for better visualization. **(F)** CCDC102B^+^ PSMC in benign-adjacent tissue co-express AR. **(G)** Small CD31^+^ vessels lined with a single layer of pericytes are demarcated by MCAM^+^CCDC102B^+^ co-immunoreactivity in high-grade PCa. The MCAM signal was omitted in the enlarged image for better visualization. **(H)** Co-expression of CCDC102B and CNN1 in PSMC in benign-adjacent (*left*) and malignant (*right)* tissues. In benign tissues, PSMC exhibit prominent membrane MCAM immunoreactivity, display a compact morphology and are arranged into densely packed bundles. In malignant tissues, dispersed and disorganised PSMC display a loss of MCAM/CCDC102B/CNN1 immunopositivity particularly in the interglandular stroma (*). Residual intact PSMC bundles are visible at the apex of tumour glands. **(I)** Pericytes encircling small vessels (yellow arrowheads) in PCa co-express MCAM and CCDC102B but lack CNN1 expression in contrast to a larger vessel (white arrowhead) with a thicker wall of MCAM^+^CCDC102B^+^CNN1^+^ VSMC. **(E-I)** Original magnification 20x. Images are representative of at least five independent experiments using tissue sections from five different patients. Source data for panels (B-D) are provided in the Source Data file.

M3 exhibited lower expression of contractile markers than M1/M2 but upregulation of synthetic genes (*HGF, MMP2, S100A10, CCL2, STEAP4* and *CCDC102B*) and gene sets related to muscle cell proliferation, connective tissue development and regulation of IGF transport/uptake (Fig. 2A-B, Supp. Fig. 1G, Supp. Table 8). M3 also expressed the highest levels of *CTGF* and *AR*, whose expression in PSMC is well-established (Brunner et al., 2025; Yang et al., 2005) (Fig. 2A, Supp. Fig. 1G). Accordingly, immunofluorescent staining of the benign prostatic stroma revealed an abundance of PLN^−^ cells in compact bundles co-expressing AR, CNN1, CCDC102B and MCAM, the latter however at lower levels than VSMC (Fig. 2E-F, 2H).

Compared to other M subclusters, M4 expressed low levels of contractile genes yet upregulation of immuno-modulatory genes and pathways (Fig. 2A-B, Supp. Fig. 1F and I), hallmarks associated with SMC dedifferentiation to a synthetic phenotype (Zhao et al., 2025). Supportively, the M4 subcluster was enriched in activated vs. non-activated tissue samples and represented in independent scRNA-seq PCa datasets (Fig. 1E, Fig. 2C, Supp. Fig. 1I). Such dedifferentiated PSMC (dPSMC) were apparent within strands of stromal tissue between tumour glands with disorganised isolated PSMC displaying decreased CNN1, CCDC102B and MCAM immunopositivity (Fig. 2G-I, Supp. Fig. 7A). Residual intact PSMC bundles typically localised to the interglandular stromal tissue at the apex of tumour acini and tumour periphery (Fig. 2H-I).

M5 expressed pericyte-associated genes (*RGS5, PDGFRB, NES, THY1, KCNJ8*) and showed strong similarity to curated pericyte signatures, including pan-cancer C8_RGS5 pericytes, healthy human prostate pericytes and pericytes from lung cancer (Chen et al., 2022; Joseph et al., 2021; Kim et al., 2020; Qian et al., 2020) (Fig. 2A and C, Supp. Fig. 1E, G and I). Consistently, CCDC102B, which was mostly strongly expressed in subcluster M5 (Fig. 2A), was co-expressed in MCAM^+^/CNN1^−^ cells that formed a single layer around small CD31^+^ vessels in both benign-adjacent and high-grade PCa tissues (Fig. 2G, 2I and data not shown). This contrasted with CNN1 expression in MCAM^+^/CCDC102B^+^ VSMC of larger vessels (Fig. 2I). Notably, the M5 gene signature correlated with multiple indicators of poor outcome, including DFS (HR 2.1; *P*=0.0086) (Fig. 2D, Supp. Fig. 3A-D).

In summary, scRNA-seq and immunofluorescent staining of clinical specimens identified multiple fibroblast and SMC/mural cell subpopulations within the benign and malignant prostate providing a framework of fibromuscular cell heterogeneity in the prostatic TME.

### Optimised tissue dissociation enhances cell yield and viability

While fibromuscular cell subpopulations identified in the re-analysed scRNA-seq dataset could be validated via staining for subcluster-enriched markers, fibroblast numbers in this dataset were low (F1-F5: 251 cells; M1-M5: 1116 cells) precluding detailed trajectory and transcriptomic analyses. This limitation is a frequent confounding issue in scRNA-seq datasets (Aparicio et al., 2025) and likely stems from suboptimal tissue dissociation of the desmoplastic stroma.

We therefore aimed to optimise tissue dissociation for enhanced recovery of fibromuscular cell subpopulations without sacrificing immune or cancer cell isolation. We first tested the ability of two commercial tissue dissociation kits (Miltenyi Biotec Tumor Dissociation Kit Human and BD Horizon Dri Tumor & Tissue Dissociation Reagent) to isolate viable stromal cell populations from a 4 mm prostate tissue punch, which represented the tissue source available herein, while maintaining an intact surface marker repertoire for flow cytometry (FC) analysis (Fig. 3A).

**Figure 3.**
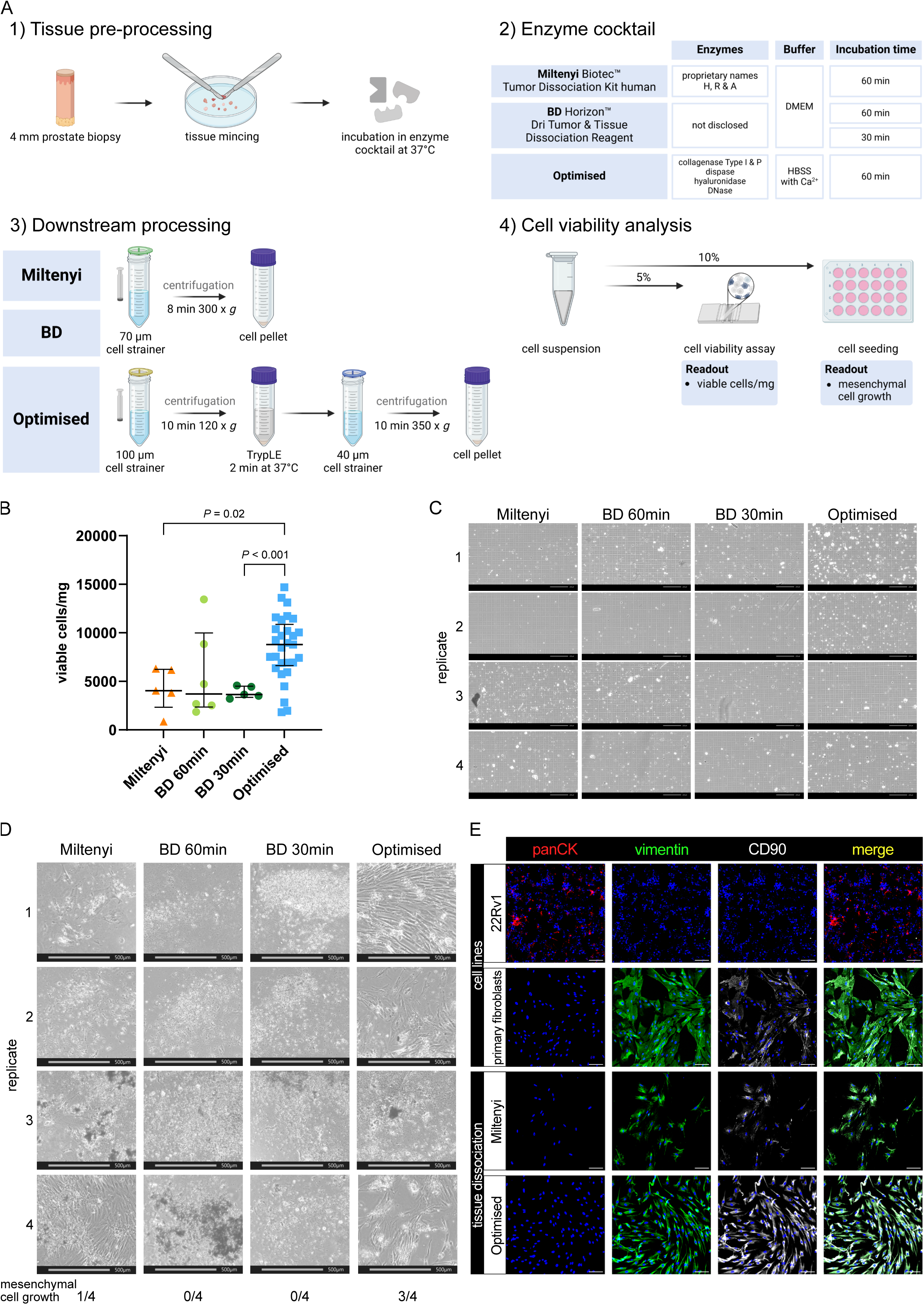
Viable cell yields differ widely between distinct tissue dissociation protocols. **(A)** Schematic comparison of the tissue dissociation protocols employed demonstrating **(1)** common tissue pre-processing steps, **(2)** enzyme cocktail composition and incubation times, **(3)** post-enzymatic dissociation processing steps and **(4)** quality assessment via cell viability analysis per trypan blue staining and cell seeding. Readouts are exemplified in B-D. **(B)** Viable cell yield/mg tissue for each protocol. Bars represent the median ± interquartile range from 5, 6 or 29 biological replicates for the Miltenyi and BD 30 min, BD 60 min or optimised protocols, respectively. Statistical significance was determined using Brown-Forsythe and Welch ANOVA tests with Dunnett T3 correction for multiple comparisons. **(C)** Brightfield imaging for trypan blue viability assessment of freshly dissociated cells isolated using the indicated protocol. Scale bars represent 200 µm at 10x magnification. **(D)** Brightfield images of the 10% seeded cells after 4 weeks. Scale bars represent 500 µm at 4x magnification. Frequency of successful mesenchymal cell growth for each of the four replicates is indicated. **(E)** Immunofluorescent staining of successfully cultured cells for the epithelial marker pan-cytokeratin (panCK) and mesenchymal markers vimentin and CD90, whereby font colour denotes pseudocolouring in the displayed images. Nuclei were counterstained using Hoechst 33342 (blue). 22Rv1 prostate cancer cells and explant cultures of primary human prostate fibroblasts served as negative and positive controls for panCK or vimentin and CD90, respectively. Scale bars represent 100 µm at 20x magnification.

The Miltenyi Biotec Tumor Dissociation Kit Human is designed for isolating diverse cell types from a broad range of tumours. Given our sample size (20–50 mg), we used a scaled-down version as recommended by Miltenyi and performed enzymatic digestion for up to 1 hour as the recommended use of a tissue dissociator was impractical for such small tissue samples. The BD Horizon Dri Tumor & Tissue Dissociation Reagent is formulated to dissociate up to 1 g of tumour tissue. To maintain an appropriate tissue-to-reagent ratio, a single vial was split between two samples permitting comparison of the recommended 30-minute incubation with an extended 60-minute incubation, the maximal duration employed for the other dissociation protocols.

Analyses revealed that both kits yielded less than 5000 viable cells/mg tissue (median: Miltenyi 4048.59 cells/mg; BD 60 min 3705.355 cells/mg; BD 30 min: 3637.4 cells/mg, Fig. 3B-C), whereby the increased digestion duration in the BD reagent did not significantly alter viable cell yields compared to the recommended 30 min duration. Notably, after four weeks of cultivating the dissociated cell suspension, only one out of four Miltenyi replicates exhibited mesenchymal cell growth, and no viable cultures were obtained using the BD reagent irrespective of digestion duration (Fig. 3D).

Due to the low yield of viable stromal cells obtained with commercial kits, we developed an optimised tissue dissociation protocol (henceforth “Optimised”) to maximise recovery of viable fibromuscular stromal cells from small prostate tissue samples. First we evaluated different buffer conditions, finding that HBSS outperformed DMEM with regards to viable cell yield (Supp. Fig. 4A), whereby all enzyme cocktails were supplemented with CaCl2 to a final concentration of 5 mM to ensure optimal activity of Ca^2+^-dependent dissociation enzymes.

Given the collagen-rich ECM of prostate tissue, we next tested various enzymes for their ability to efficiently dissociate tissue samples without sacrificing cell viability, observing that 2 mg/ml Collagenase Type 1 improved viable cell yields per mg tissue compared to 4 mg/ml (Supp. Fig. 4A). To ensure consistency across enzyme batches, we calculated the enzyme activity of the 2 mg/ml Collagenase Type I and established 546 U/ml as the standard activity for all subsequent experiments. The addition of 1.5 U/ml Collagenase P further improved viable cell yield (Supp. Fig. 4A). Besides collagen, tissue ECM comprises proteoglycans like hyaluronan (Reichard and Asosingh, 2019) with hyaluronidase a common component of tissue dissociation reagents (Costa et al., 2018). However, compared to the concentrations typically employed (e.g. 2 mg/ml in (Costa et al., 2018)), lower amounts of hyaluronidase (100 µg/ml) were sufficient to improve tissue dissociation herein (Supp. Fig. 4A). Additionally, 0.6 U/ml Dispase II, a neutral protease targeting fibronectin and collagen IV (Reichard and Asosingh, 2019), was included to facilitate ECM degradation.

During tissue dissociation, free DNA released by dead cells leads to cell/tissue aggregation, which can be overcome by incorporating DNase I into the dissociation cocktail (Reichard and Asosingh, 2019; Slyper et al., 2020). We initially used DNase I at a concentration of 25 µg/ml as previously reported (Costa et al., 2018; Quatromoni et al., 2015), data not shown) but found that increasing it to 100 µg/ml as per (Dominguez et al., 2020; Slyper et al., 2020) prevented the minced tissue from clumping, presumably thereby increasing the surface area available for enzymatic digestion.

For the 37°C incubation of the tissue in the enzyme cocktail, we opted for a waterbath rather than an incubator to allow real-time visual monitoring without temperature fluctuations. Prolonged incubation in enzymatic cocktails negatively impacts cell viability (Reichard and Asosingh, 2019). We observed considerable inter-sample heterogeneity with some tissue samples more readily dissociating than others. Thus, to avoid over-digestion of readily dissociated samples and under-digestion of more resistant samples, reactions were terminated either upon the lack of macroscopically-visible tissue pieces or upon a maximum of 1 hour, whichever was attained first.

Compared to the commercial kits, the optimised protocol incorporated additional processing steps to improve single-cell recovery of cell clusters potentially lost in straining steps. Hereby, the dissociation mixture was first strained through a 100 µm cell strainer, then centrifuged at 120 × g for 10 min at 4°C, a gentler condition compared to commercial protocols. The resulting pellet was resuspended in the trypsin-like protease TrypLE and incubated at 37°C for 2 minutes to dissociate cell-cell contacts. A second straining step through a 40 µm filter was followed by final centrifugation at 350 × g for 10 min at 4°C, optimising recovery of a homogeneous single-cell suspension before final resuspension in 100 µl of FC buffer.

Collectively, these optimisation steps more than doubled the viable cell yield compared to commercial kits, reaching a median of 8788.07 cells/mg tissue (Fig. 3B). Additionally, the optimised protocol tended to recover a greater proportion of cells >10 µm in diameter, while the proportions of cells with diameters <5 µm and between 5-10 µm remained comparable across protocols (Supp. Fig. 4B). This suggested enhanced isolation of larger cells, such as stromal, epithelial and endothelial cells, relative to smaller immune cells.

Importantly, 3 of 4 samples dissociated using the optimised protocol resulted in stromal cell growth (Fig. 3D). Since we ultimately aimed to better characterise the fibromuscular component of the prostate TME, we confirmed the mesenchymal origin of cells that, following tissue dissociation, could be cultured/passaged under conditions that support propagation of fibroblasts but not epithelial or endothelial cells (Fig. 3E).

In summary, the optimised tissue dissociation protocol significantly increased cell yield and viability compared to the commercial kits tested, and enabled successful isolation and culture of mesenchymal cells from small prostate biopsy samples.

### Optimised tissue dissociation significantly increases the yield of CD31^−^ stroma

Tissue-dissociated single cell suspensions were subsequently analysed via FC to (1) monitor the relative distribution of dead cells, debris vs. (single) viable cells across the different protocols, (2) identify the broad cell types recovered, and (3) assess the number of stromal cells isolated (Fig. 4A). Thus, the 85% of cell suspension remaining after the aforementioned quality controls (Fig. 3A, panel 4) was stained with an FC marker panel to identify leukocytes (CD45), basal and non-basal prostate epithelial cells (CD326^+^PDPN^+^ and CD326^+^PDPN^−^, respectively), blood endothelial cells (BEC) and lymphatic endothelial cells (LEC) (CD31^+^PDPN^−^ vs. CD31^+^PDPN^+^, respectively). Seven stromal-associated cell surface markers (MCAM, CD140a, CD140b, CD90, PDPN, FAP, CD105) were additionally included in the 11-channel panel to discriminate potential stromal subpopulations in the remaining cells (hereafter termed CD31^−^ stroma), and were selected based on literature and their differential expression across the scRNA-seq-derived F/M subclusters (Supp. Fig. 1G, Fig. 6P and Supp. Table 11). Of note, CD140a, CD140b, CD90 and CD105 are encoded by *PDGFRA, PDGFRB, THY1* and *ENG,* respectively. A gating strategy (Fig. 4B) was applied to single viable cells permitting the stepwise exclusion of immune/endothelial/epithelial cells until only the CD31^−^ stroma cells of primary interest to the current study remained. The CD31^−^ stromal fraction was subsequently analysed further by applying single gates to determine the proportion of cells positive for each stromal cell surface marker (Fig. 4B viii). Importantly, signal intensities for these different markers were highly comparable across the different protocols (Supp. Fig. 4C) indicating that the dissociation protocols employed did not diverge with respect to loss/over-digestion of these markers.

**Figure 4.**
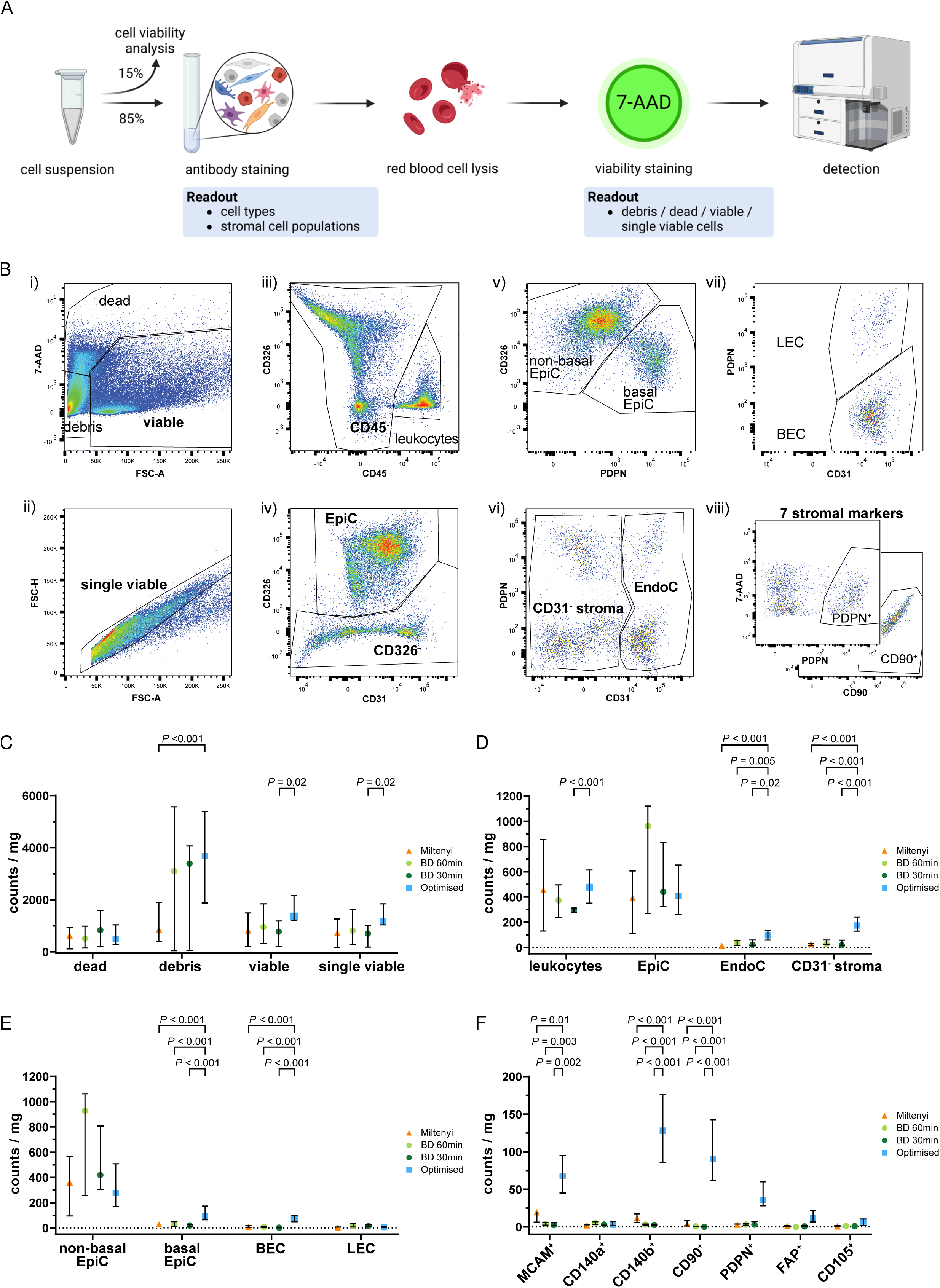
Flow cytometry analysis of single cell suspensions after tissue dissociation. Schematic representation of FC staining of the single cell suspension remaining after tissue dissociation and cell seeding/viability analyses. **(B)** Representative gating strategy used to sequentially identify different cell types whereby **(i)** staining with 7-AAD identified viable cells, which were used to **(ii)** gate for single viable cells based on FSC-A and FSC-H. Consecutive exclusion of **(iii)** leukocytes (CD45^+^), **(iv)** epithelial cells (CD326^+^; EpiC), **(v)** basal epithelial (CD326^+^PDPN^+^) or non-basal epithelial (CD326^+^PDPN^−^) cells, **(vi)** endothelial cells (CD31^+^; EndoC), **(vii)** blood endothelial cells (CD31^+^PDPN^−^; BEC) or lymphatic endothelial cells (CD31^+^PDPN^+^; LEC) resulted in the remaining CD31^−^ stroma, which was **(viii)** subsequently used for single gating of seven different stromal markers (MCAM, CD140a, CD140b, CD90, PDPN, FAP, CD105). Representative plots for PDPN and CD90 are shown. **(C-F)** Flow cytometry analysis for each tissue dissociation protocol showing in counts/mg tissue the percentages of **(C)** dead, debris, viable and single viable cells, **(D)** leukocytes, EpiC, EndoC, and CD31^−^ stroma, **(E)** EpiC and EndoC further distinguished by PDPN expression into non-basal or basal EpiC, and BEC or LEC, **(F)** CD31⁻ stromal cells positive for each of the stromal markers. **(C-F)** Bars represent the median ± interquartile range. Statistical significance was determined using **(C-E)** mixed-effects model with the Geisser-Greenhouse correction and Dunnett correction for multiple comparisons or **(F)** 2-way ANOVA with Dunnett’s correction for multiple comparisons. **(C-F)** Data are derived from multiple independent experiments using tissue samples from different patients whereby n = 4-6 (Miltenyi), 3-6 (BD 60min and 30 min) or 27-29 (Optimised).

The optimised protocol showed a trend towards the lowest percentage of dead cells but yielded significantly more debris than Miltenyi samples, possibly indicating enhanced ECM dissociation, although debris counts/mg tissue were comparable between the BD samples and the optimised protocol (Fig. 4C). Consistent with the significantly higher number of viable cells/mg tissue using the optimised protocol (Fig. 3B), these samples tended to yield the highest number of (single) viable cells/mg tissue with a significant increase over BD 30 min samples (Fig. 4C).

Leukocyte and epithelial cell (EpiC) counts were comparable across protocols. Compared to commercial kits however, the optimised protocol significantly increased EndoC and CD31^−^ stromal cell counts/mg tissue also when calculated as proportions of single viable cells (Fig. 4D, Supp. Fig. 4E, Supp. Table 12). Considerable variation in the ratio of non-basal (luminal and intermediate) to basal EpiC was observed between protocols, whereby >90% of EpiC isolated by commercial kits were non-basal whereas the optimised protocol recovered 27.9% basal EpiC (Supp. Fig. 4G, Supp. Table 12). This enhanced isolation of basal EpiC was also evident when analysing counts/mg tissue and as a proportion of single viable cells (Fig. 4E, Supp. Fig. 4F). Similarly, the optimised protocol yielded a significantly higher percentage of BEC compared to the BD kit (Supp. Fig. 4G) but at similar proportions to the Miltenyi kit. Consequently, LEC comprised only 10.8% of EndoC isolated by the optimised protocol but represented the dominant EndoC type isolated by the BD kit (Supp. Fig. 4G, Supp. Table 12). When considering absolute counts/mg tissue or proportions of single viable cells however, the optimised protocol yielded significantly higher numbers of BEC while LEC proportions were comparable across protocols (Fig. 4E and Supp. Fig. 4F, Supp. Table 12).

In summary, the optimised protocol demonstrated superior isolation of tissue-resident cell types, such as basal EpiC, BEC and CD31^−^ stromal cells over the commercial kits tested, suggesting greater tissue dissociation and consequently improved representation of cellular heterogeneity.

### Optimised tissue dissociation enhances isolation of cells expressing diverse fibromuscular markers

The CD31^−^ stromal fraction encompassed the cell populations of primary interest to the current study. To assess the ability of the optimised protocol to isolate distinct fibromuscular cell subtypes, we calculated the percentage of cells expressing each stromal marker within the CD31⁻ stromal compartment. This provided insight into the relative abundance of specific marker-positive subpopulations but failed to account for variations in the total yield of CD31⁻ stromal cells across protocols (Fig. 4D and Supp. Fig. 4I). Thus, to more accurately compare the different protocols, we calculated the absolute counts of marker-positive cells/mg tissue, as well as their proportion among single viable cells. Theses analyses revealed the optimised protocol yielded superior (MCAM, CD140b, CD90, PDPN, FAP, and CD105) or comparable (CD140a) numbers of cells expressing these markers relative to the commercial kits tested (Fig. 4F, Supp. Fig. 4H), implying that the optimised dissociation protocol significantly improves recovery of heterogeneous stromal cell types.

### Optimised dissociation of benign and malignant prostate tissue yields comparable amounts of viable stromal cells

Optimisation experiments thus far employed benign tissue to minimise variations in sample composition, which could confound comparison of dissociation protocols. We therefore next evaluated the efficacy of the optimised protocol to dissociate biopsy cores from macroscopically-suspected malignant and patient-matched benign-adjacent tissues. Histopathological validation (Fig. 5A and Supp. Fig. 5, Supp. Table 14) identified eight patients for whom a tumour-containing biopsy core and ≥1 matching benign core were dissociated. This hereon termed eight patient-matched cohort comprised eight cancer tissue (CA) samples and 12 matched benign-adjacent (BE) samples. Biopsy cores from the remaining patients (n=10) contained only benign tissue and were employed to increase depth of the benign dataset (termed hereafter benign-enriched cohort).

**Figure 5.**
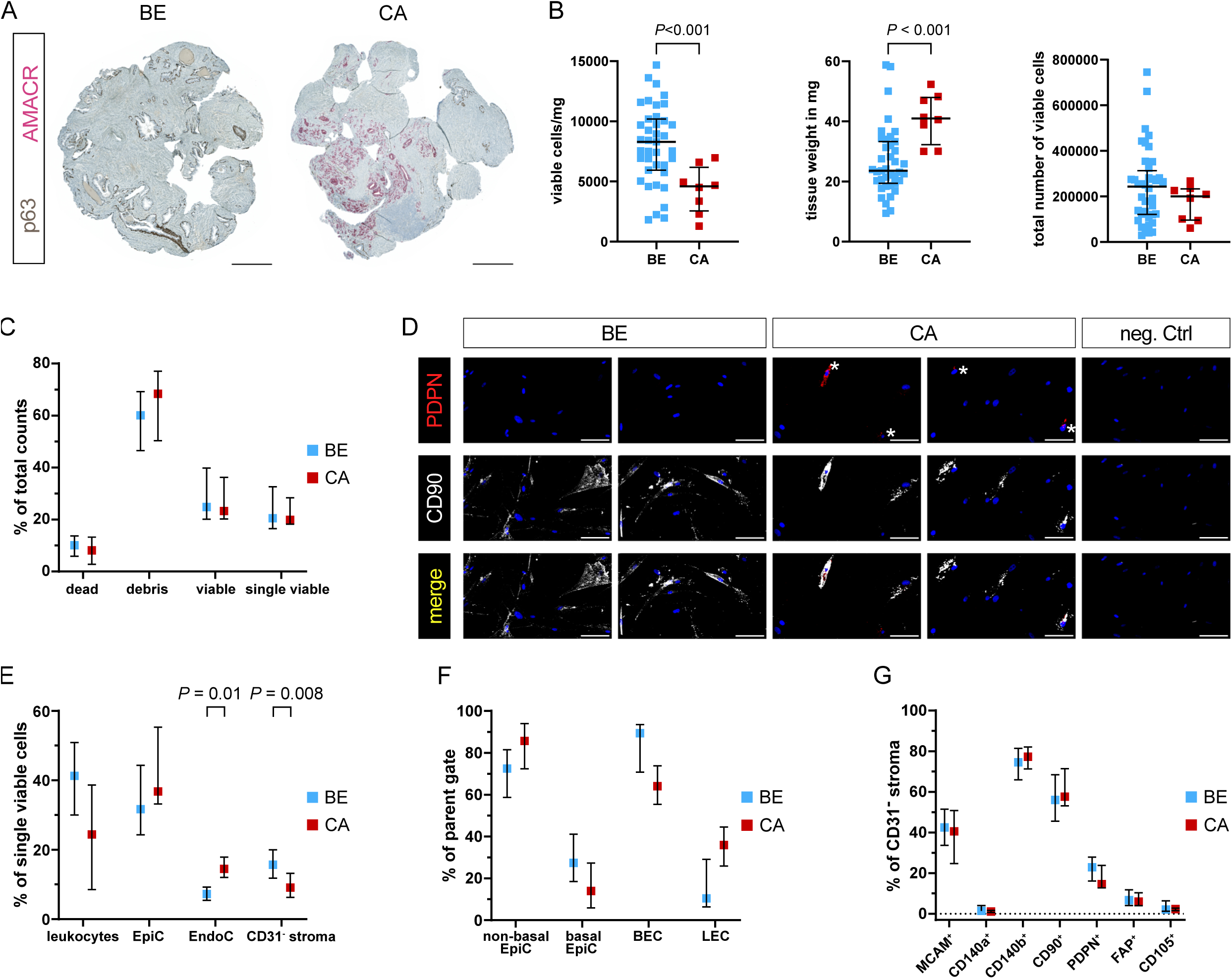
Enhanced dissociation of malignant and benign-adjacent prostate tissues reveals disease-associated differences in cellular composition. **(A)** Dual immunohistochemistry of the basal epithelial marker p63 (brown) and tumour cell marker AMACR (red) aiding histopathological assessment of 4 mm biopsy cores sampled from benign (BE) or cancerous (CA) tissue regions. Representative images from patient 4 are shown. Scale bars represent 500 µm at 20x magnification. **(B-G)** Readouts of tissues from the eight patient-matched and benign-enriched cohorts dissociated using the optimised protocol. **(B)** Viable cell yield/mg tissue *(left)*, tissue weight of biopsy cores in mg *(centre)* and total viable cell number *(right)*. Bars represent the median ± interquartile range from ≥39 benign (BE) and eight tumour (CA) replicates. **(C)** FC analysis showing dead, debris, viable and single viable cells as percentages of total counts. **(D)** Immunofluorescent staining of the stromal cell markers PDPN (* denotes positive cells) and CD90 in cells successfully cultured from benign and tumour-containing tissue-dissociated samples, whereby font colour denotes pseudocolouring in the images displayed. Nuclei were counterstained using Hoechst 33342 (blue). Scale bars represent 100 µm at 20x magnification. Negative control (neg. Ctrl, *right*) incubated without primary antibodies is shown. **(E-G)** FC analysis depicting **(E)** leukocytes, epithelial cells (EpiC), endothelial cells (EndoC), and CD31^−^ stroma as a percentage of single viable cells; **(F**) EpiC and EndoC further distinguished by PDPN expression into non-basal or basal EpiC, and blood endothelial cells (BEC) or lymphatic endothelial cells (LEC), as a percentage of their respective parent gate (EpiC or EndoC); **(G)** cells positive for each stromal marker expressed as a percentage of the CD31^−^ stroma. **(C, E-G)** Bars represent the median ± interquartile range. Data are derived from multiple independent experiments using tissue samples from different patients whereby n=36-39 benign and 5-8 cancer samples. Statistical significance was determined using (B) Welch’s t test, (C-F) mixed-effects model with the Geisser-Greenhouse correction and Šídák correction for multiple comparisons, or (G) 2-way ANOVA with Šídák correction for multiple comparisons.

Although malignant and benign tissue biopsy cores were comparable in size, we unexpectedly observed a decrease in the number of viable cells from tumour compared to benign-adjacent cores when adjusted per mg tissue (Fig. 5B left panel, Supp. Table 13). Concomitantly however, malignant biopsy cores were significantly heavier than benign tissue cores (Fig. 5B middle panel), suggesting the decreased number of viable cells/mg from malignant tissues may arise from the increased weight of the tumour-containing cores – for example due to increased ECM density, a phenomenon also noted in the literature (Dogan et al., 2005). Indeed, the total number of viable cells was comparable between malignant and benign-adjacent tissue cores (Fig. 5B right panel) and no significant differences were observed with respect to debris or dead/viable/single viable cells (Fig. 5C). Importantly, cells isolated using the optimised protocol could be successfully cultured from both benign and tumour-containing biopsy cores and expressed mesenchymal markers (Fig. 5D).

Whilst EndoC were significantly enriched, the proportion of CD31^−^ stroma was decreased in malignant *versus* benign samples (Fig. 5E, Supp. Table 13). No significant difference was observed in the expression level of any of the stromal surface markers in the malignant/benign CD31^−^ stromal fractions (Fig. 5G), suggesting over-digestion was not the primary underlying cause for this effect. Rather, expansion of (malignant) epithelial cells in PCa tissue at the expense of the fibromuscular stroma may have contributed to the observed relative reduction in CD31^−^ stroma. This hypothesis was further supported by the trend towards higher percentages of EpiC/non-basal EpiC in malignant compared to benign-adjacent samples (Fig. 5E-F, Supp. Table 13), whereby basal EpiC loss is a key PCa diagnostic feature (Humphrey, 2017). Likewise, malignant samples displayed marked trends towards a lower proportion of leukocytes and BEC vs. LEC (Fig. 5F) potentially indicative of increased lymphatic vessel density and decreased immune cell infiltration, both established features of PCa (Datta et al., 2010; Stultz and Fong, 2021).

Collectively, data thus far indicate the optimised protocol efficiently dissociates both malignant and benign-adjacent tissue samples enabling the isolation and cultivation of mesenchymal cells that retain expression of multiple fibromuscular surface markers.

### Stromal marker clustering identifies distinct tumour- and benign-enriched populations

Since characterisation of stromal heterogeneity represented the overarching goal of this study, dimensionality reduction of CD31^−^ stroma FC counts was performed with tSNE for the eight patient-matched cohort to identify putative benign- or tumour-enriched stromal cell subpopulations (Fig. 6, Supp. Fig. 6A). While tumour and benign samples were largely homogeneously distributed across the tSNE plot and patients, two subpopulations that mainly derived from patients 1 and 3 primarily originated from malignant or benign-adjacent samples, respectively (Fig. 6A-D, Supp. Table 15).

**Figure 6.**
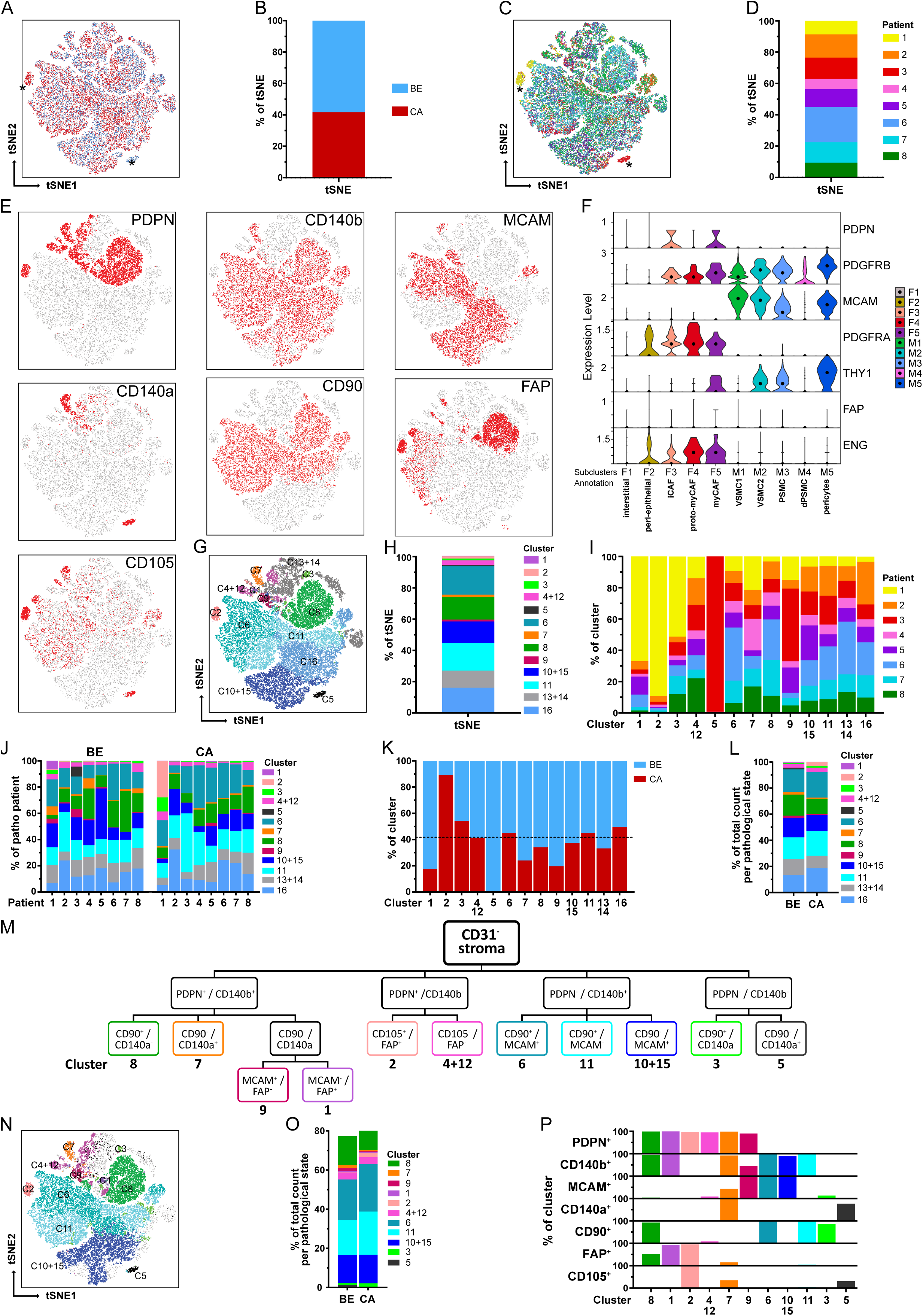
Identification of stromal clusters in benign vs. malignant prostate tissues via tSNE and stromal surface marker-based gating. **(A-H)** tSNE dimensionality reduction analysis of CD31^−^ stroma from flow cytometry data based on stromal markers. Matched benign (BE, n=12) and cancer (CA, n=8) samples from eight patients were pooled. Sample counts were stratified to visualise **(A-B)** BE (blue) vs. CA (red) sample counts, **(C-D)** counts derived per patient, **(E)** cells positive for each of the single stromal markers and **(G-H)** clusters identified by XShift post merge. **(A, C)** Asterisks mark benign- and cancer-specific populations. **(B, D, H)** Stacked bar plots depicting relative distributions as percentages of the tSNE. **(F)** Violin plot depicting mRNA levels of the seven stromal surface markers in the fibromuscular cell subclusters of the re-analysed PCa scRNA-seq dataset. **(I)** Relative distribution of patient samples across the 13 combined clusters. **(J)** Relative distribution of the 13 combined clusters in the eight patients according to the BE and CA pathology of the corresponding tissue core. **(K)** Relative distribution of BE and CA sample counts per cluster in relation to overall BE and CA sample count distribution whereby the dotted line denotes 41.7% of total sample counts that derived from CA samples. **(L)** Stacked bar plot displaying the relative distribution of the 13 combined clusters in BE vs. CA sample counts. **(M)** Gating strategy developed using stromal surface markers ranked according to increasing cluster-specific expression identifying 11 of the 13 combined clusters. **(N)** tSNE plot overlaid with the 11 gated clusters. **(O)** Stacked bar plot showing the relative distribution of gated clusters in BE vs. CA sample counts. **(P)** Expression of the seven stromal surface markers across the 11 gated clusters.

Overlaying the CD31^−^ stroma counts positive for each stromal marker on the tSNE map enabled visualisation of regions enriched for single or multiple stromal markers (Fig. 6E). However, for unbiased identification of stromal clusters independent of the tSNE plot, unsupervised clustering was performed using XShift (Samusik et al., 2016) for the prominently expressed stromal markers MCAM, CD140b, CD90, PDPN and FAP (Fig. 6E). CD140a and CD105 were excluded in this step due to overclustering (see Methods). These analyses segregated CD31^−^ stroma counts into 16 clusters (Supp. Fig. 6B-G, Supp. Table 15), whereby those expressing similar marker combinations were considered “overclustered” (Supp. Fig. 6G) and subsequently merged yielding 13 clusters with distinct stromal marker expression patterns (Fig. 6G-H, Supp. Table 15). The resulting cluster boundaries (Fig. 6G) closely overlapped with regions positive for single-marker expression on the tSNE plot (Fig. 6E). Moreover, median fluorescence intensity values for each stromal marker differed substantially across the XShift clusters (Supp. Table 16) further supporting the robustness of this clustering approach. Six clusters (clusters 6, 8, 11, 16 and the merged clusters 10+15 and 13+14) constituted the majority (90.6%) of the tSNE plot and were distributed similarly across patients (Fig. 6H). With the exception of patient 3-specific cluster 5, all other clusters were present across all patients, although we noted the patient 1 CA sample exhibited a markedly different cluster distribution compared to the other CA samples (Fig. 6I-J).

Since the tSNE plot comprised 58.3% BE vs. 41.7% CA sample counts, we reasoned that clusters originating equally from benign and malignant samples would be expected to show a similar distribution (Fig. 6K), whereas clusters deviating considerably from this distribution may indicate enrichment for either benign- or tumour-enriched samples. Supportively, clusters 2 and 5 showed strong enrichment in malignant and benign samples, respectively, and represented the clusters assigned by XShift to the aforementioned tumour- and benign-enriched populations from patients 1 and 3 (Fig. 6A, C, G and K). Moreover, this approach identified further clusters potentially enriched in benign (clusters 1, 7 and 9) or malignant (cluster 3) samples (Fig. 6K).

### A stromal marker-based gating strategy discriminates 11 clusters encompassing distinct fibromuscular cell types

To identify surface marker combinations capable of discriminating these stromal subpopulations for their future isolation, a gating strategy was developed by ranking markers expressed in the 13 combined clusters according to their ability to distinguish broader groups (highest ranking) *vs.* more specific clusters (lowest ranking; Supp. Fig. 6I, 6K, 6M). Applying this strategy to the CD31^−^ stroma of the eight patient-matched cohort enabled identification of 11 of 13 clusters based on its unique surface marker profile representing >75% of the benign and malignant CD31^−^ stroma (Fig. 6O, Supp. Table 15). Cluster 13+14 could not be positively selected since it lacked expression of all panel markers and cluster 16 split upon applying the gating strategy such that its FC counts were reassigned to clusters 6, 11, and 10+15 (Fig. 6N) most likely due to the stricter positive marker cut-offs employed in the gating strategy compared to the interpretation of ‘positive’ and ‘negative’ signals by XShift.

To annotate FC-gated stromal subpopulations (Fig. 6N), we compared their surface marker profiles to the fibromuscular clusters of the re-analysed scRNA-seq dataset (Fig. 6F, P) and undertook immunofluorescent staining of prostate tissue specimens (Fig. 7-8 and Supp. Fig. 7). Clusters 3 and 13+14 could not be unequivocally identified due to the paucity of markers expressed by these clusters (Fig. 6P and Supp. Fig. 6I). The patient 3-specific cluster 5 expressed BEC markers CD140a and CD105 but lacked expression of the other FC panel stromal markers and was thus considered to represent contaminating endothelial cells potentially due to diminished CD31 expression, an established event in inflammation (Cheung et al., 2020; Kato et al., 2019) and which was readily apparent together with PIN in patient 3 tissue cores (Supp. Table 14, Fig. 6P).

**Figure 7.**
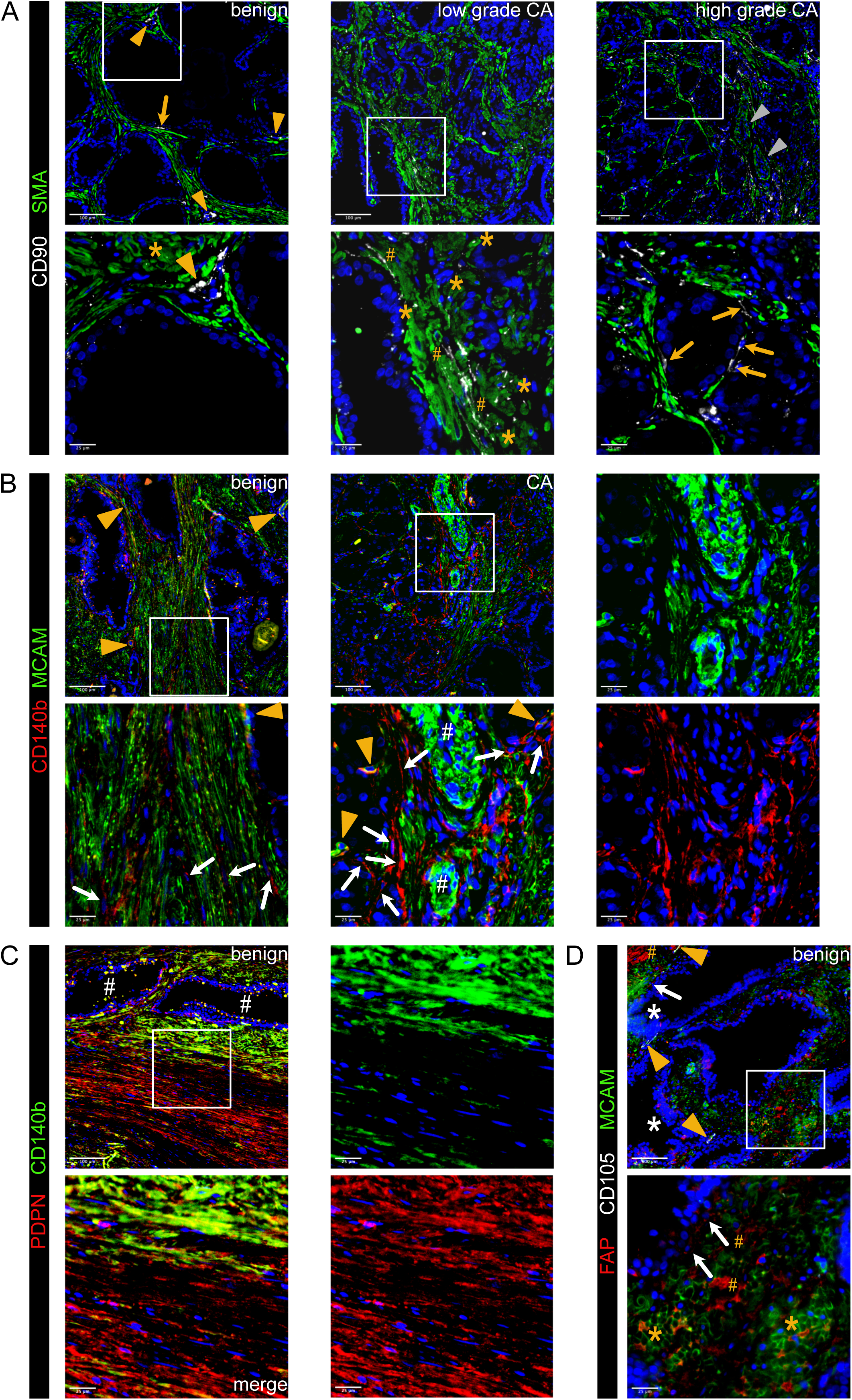
Immunofluorescent staining of prostate tissues using flow cytometry stromal panel markers identifies distinct subpopulations of fibromuscular stromal cells (A-D) Immunofluorescent staining of the antibodies stated in prostate tissue sections of indicated pathology. Boxed regions are enlarged beneath the parental image. Font colour corresponds with the pseudocolouring of the antigen stained. Images are representative of at least three independent experiments using tissues derived from at least five patients. **(A)** CD90^+^SMA^+^ cells surround small vessels (orange arrowheads, *left*) whereas CD90^−^SMA^+^ cells surround large vessels (grey arrowheads, *right*). Peri-epithelial CD90^+^SMA^−^ cells with elongated fibroblast-like morphology (orange arrows, *top left and bottom right*), and elongated CD90^+^ cells adjacent to SMA^+^ SMC bundles (orange hashtags) are highlighted. Asterisks demarcate CD90^+^SMA^+^ cells within intact SMC bundles. **(B)** Co-expression of MCAM and CD140b in PSMC of benign and malignant tissues with extensive co-localisation in the wall of small vessels consistent with pericytes (orange arrowheads) but not in multi-layered wall of larger vessels (white hashtags). CD140b^+^MCAM^−^ cells with an elongated fibroblast-like morphology (white arrows) are observed particularly in peri-tumoral regions but also interspersed between SMC bundles of benign tissues. **(C)** A single PDPN^+^ layer of basal epithelial cells is indicative of benign glands (white hashtags). Example of a focal area exhibiting intense stromal PDPN staining, whereby the glands are encircled by a layer of PDPN^+^CD140b^+^ PSMC juxtaposed to a distinct more distal layer of PDPN^+^CD140b^low^ PSMC. **(D)** Focal areas of benign tissue exhibiting co-expression of FAP^+^MCAM^+^ PSMC (asterisks) interspersed with FAP^+^MCAM^−^ cells (hashtags). Heterogeneous FAP expression by some epithelial cells is further suggestive of tissue activation with some glands exhibiting morphological features associated with PIN (white asterisks). Orange arrowheads demarcate blood vessels with CD105^+^ endothelial cells.

**Figure 8.**
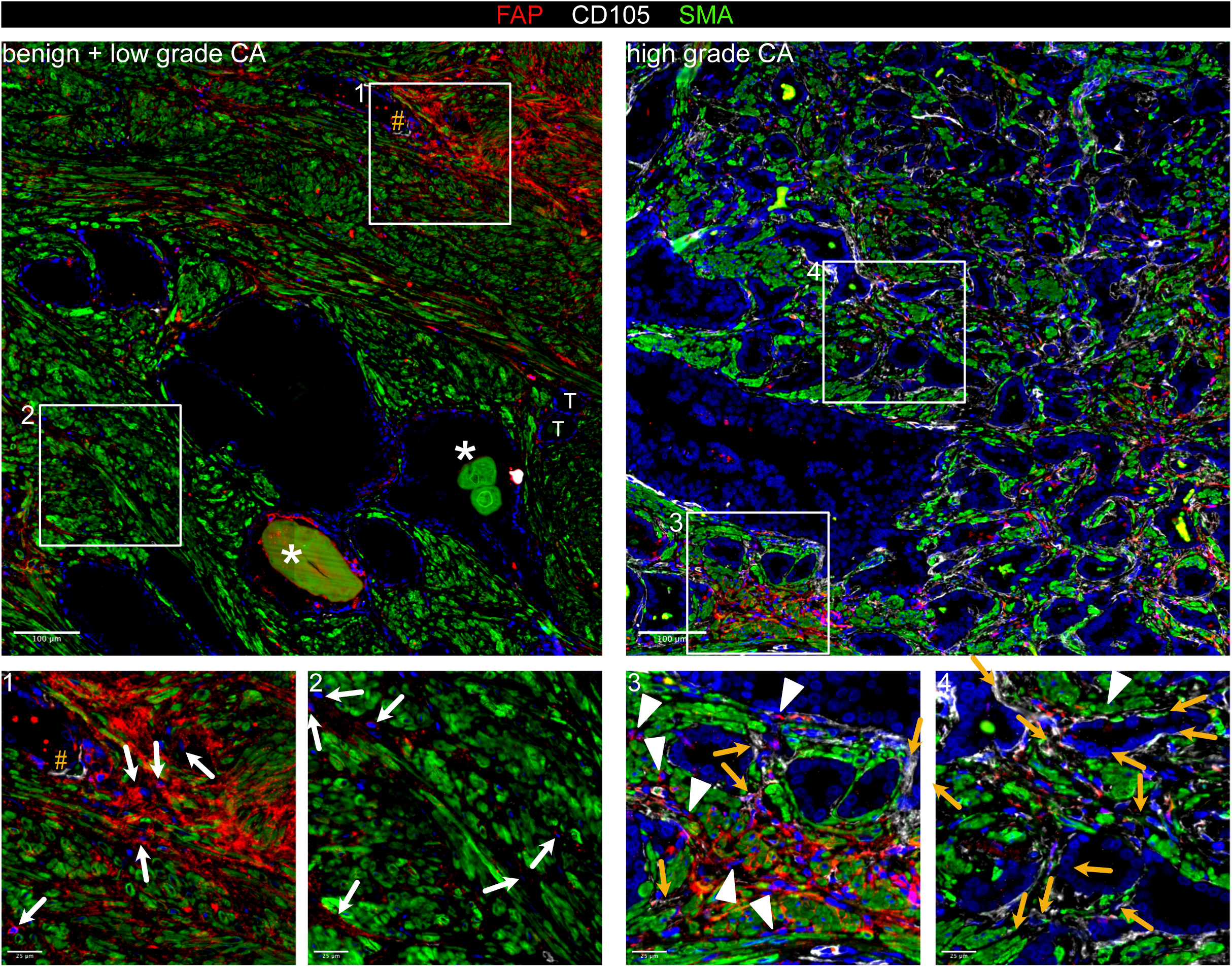
Co-expression of FAP and CD105 demarcate myCAF in prostate cancer. Immunofluorescent staining of FAP, CD105 and SMA in prostate tissue sections of indicated pathology. Boxed regions are shown enlarged beneath the parental image. Font colour corresponds with the pseudocolouring of the antigen stained. Images are representative of at least three independent experiments using tissues derived from at least five patients. Enlarged region 1 demarcating focal stromal activation with extensive FAP expression by SMA^+^ PSMC but also in interspersed SMA^−^ cells (white arrows), which are also identified in enlarged region 2. Orange hashtags denote blood vessels with CD105^+^ endothelial cells. Asterisk marks benign glands with corpora amylacea. Tumour glands are labelled (T). Enlarged regions 3 and 4 displaying an abundance of CD105^+^FAP^+^SMA^+^ myCAF (orange arrows) with an elongated fibroblast morphology and primarily localised adjacent to tumourigenic glands. Interspersed CD105^−^FAP^+^SMA^−^ cells (presumptive iCAF) are highlighted (white arrowheads).

On the basis of their MCAM^+^ profile, clusters 6, 9, 10+15 were considered to represent distinct SMC/mural cell subpopulations, whereby the CD90^−^ status of cluster 10+15 aligned with that of VSMC1, whereas the CD90^+^ cluster 6 was similar to VSMC2, PSMC and pericytes (Muhl et al., 2020) (Fig. 2E, H-I and 6F, M, P). Supportively, CD90^+^SMA^+^ cells were observed in the wall of small vessels but not in the multi-layered wall of larger vessels (Fig. 7A) with SMA immunostaining employed to identify SMC due to decreased MCAM immunofluorescence of PSMC in PCa tissues (Fig. 2E, H and Supp. Fig. 7A). Partial co-expression of CD90 in SMA^+^ PSMC, which co-expressed MCAM and CD140b (Fig. 7A-B), further supported annotation of cluster 6 as a mix of VSMC2, PSMC and pericytes. The subset of FAP^+^ cells within cluster 6 (Fig. 6E) were considered to denote activated PSMC, which were observed in nests at the tumour front and in activated benign-adjacent tissues, particularly adjacent to glands exhibiting PIN, basal cell hyperplasia or corpora amylacea (Fig. 7D and Fig. 8). The benign-enriched cluster 9 co-expressed PDPN, MCAM and CD140b (Fig. 6K, M, P). While basal epithelial cells and lymphatic vessels express PDPN, these cells lack MCAM and CD140b. Rather, the PDPN^+^MCAM^+^CD140b^+^ profile of cluster 9 was strikingly similar to PSMC within the fibrotic stroma of benign hyperplastic nodules (Fig. 7C and Supp. Fig. 7B-C).

Cluster 11 displayed a PDPN^−^/CD140a^−^/FAP^−^/MCAM^−^/CD140b^+^/CD90^+^ profile and was abundant in both benign and malignant samples (Fig. 6J-P). Beyond the aforementioned expression of CD90^+^SMA^+^cells in small vessels and PSMC (cluster 6, Fig. 6P and 7A), CD90 was also detected in SMA^−^ cells exhibiting an elongated fibroblast-like morphology adjacent to epithelial glands and along SMC bundles (Fig. 7A) similar to previously reported “wrapping” interstitial fibroblasts (Joseph et al., 2021; Peng et al., 2013). A similar distribution was also observed for CD140b^+^/MCAM^−^ cells in benign tissues (Fig. 7B). Cluster 11 was therefore considered to represent non-activated fibroblasts.

Cluster 7 constituted a minor subpopulation enriched in the benign cores from patients 1 and 4 but was also present in malignant samples and broadly expressed CD140a, PDPN and CD140b (Fig. 6I-P). A subset of cells additionally expressed MCAM (Fig. 6P). Whilst the MCAM^−^ fraction (57.2% of cluster 7) largely lacked FAP and CD105 expression consistent with an early-activated fibroblast phenotype, the MCAM^+^ fraction (42.9% of cluster 7) displayed a greater abundance of FAP^+^ cells (23.4%), >90% of which co-expressed CD105 (Fig. 6E, P and Supp. Fig. 6J). Immunofluorescent staining revealed the presence of MCAM^+^FAP^+^CD105^+^ cells surrounding small vessels lacking a multi-layered vascular wall (Supp. Fig. 7E) implying cluster 7 may harbour a pericyte subset consistent with reports that CD140a^+^CD140b^+^CD34^+^ pericytes can express PDPN, FAP and CD105 under inflammatory and tumourigenic conditions (Cimini and Kishore, 2021; Ebert et al., 2020; Rivera and Brekken, 2011). The MCAM^+^FAP^−^ fraction (74.7%) comprised both CD105-positive and -negative cells (30.0% and 44.7%, respectively) whose identity could not be discerned with confidence. It may be noted however that some non-vessel-associated FAP^−^CD105^+^ cells expressed the PSMC/mural cell marker MCAM (Supp. Fig. 7D) potentially suggestive of SMC-/pericyte-derived myCAF or myofibroblasts as previously described (Gao et al., 2024; Hosaka et al., 2016).

The remaining gated clusters 1, 2, 8 and 4+12 were MCAM^−^/PDPN^+^ indicative of activated fibroblasts in accordance with our scRNA-seq analyses (Fig. 6F) (Henry et al., 2018). The minor subpopulation cluster 4+12 however expressed only PDPN and could not be unequivocally annotated since non-stromal PDPN^+^ entities, such as peripheral neurons, basal EpiC and LEC (Fig. 6P and Supp. Fig. 7B-C), would also be expected to be negative for other stromal FC panel markers. Cluster 1, 2 and 8 were FAP^+^ further supporting their annotation as activated fibroblast subpopulations, whereby clusters 1 and 8 also expressed CD140b but differentially expressed CD90 (Fig. 6M, P). Cluster 1 represented a minor subpopulation primarily isolated from the benign sample of patient 1 and lacked CD90 expression implying cluster 1 may denote fibroblasts at an early stage of activation (Fig. 6J, P) (True et al., 2010). In contrast, the CD90^+^ cluster 8 constituted an abundant subcluster isolated at comparable frequency from benign and malignant cores (Fig. 6H, J-L, O-P) and exhibited a surface marker profile similar to CAF-S5, whose lack of contractile markers (SMA, TAGLN and TPM2) but expression of *C3* and *C7* and localization distal to tumour glands (Mathieson et al., 2024), were highly reminiscent of subcluster F3/iCAF (Fig. 1G, Supp. Fig. 1F-G and Supp. Fig. 2). Indeed, iCAF are present in both benign-adjacent and malignant tissues with CD90^+^ fibroblasts associated with inflammatory processes (Brunner et al., 2025; Jenkins et al., 2025; Zeng et al., 2023). Cluster 8 was thus considered to represent iCAF. In line with these annotations, both cluster 1 and 8 lacked the myCAF marker CD105 (Fig. 6P). Moreover, FAP^+^ or CD140b^+^ cells with an elongated fibroblast-like morphology and that lacked co-expression of the PSMC markers MCAM/SMA and myCAF marker CD105 were observed in benign and malignant tissues (Fig. 7B-C and Fig. 8). Robust expression of CD105 however in cluster 2 together with its PDPN^+^/FAP^+^ status and enrichment in malignant samples (Fig. 6O-P) suggested this cluster represented myCAF. Supportively, malignant tissues exhibited strong upregulation of CD105, whereby CD105^+^FAP^+^ cells primarily localised to the peri-tumoral space, the established myCAF niche (Brunner et al., 2025; Croizer et al., 2024). Furthermore, CD105^+^FAP^+^ cells co-expressing SMA but largely lacking MCAM were also readily apparent and distinguishable from residual CD105^−^FAP^+/-^SMA^+^ PSMC (Fig. 8 and Supp. Fig. 7D).

In summary, thirteen stromal clusters exhibiting distinct surface marker profiles were identified in the PCa and benign prostate microenvironment, eleven of which could be discriminated using a gating strategy. By cross-referencing the gated FC clusters with independent scRNA-seq data and immunofluorescent staining, several FC-identified clusters were mapped to distinct parenchymal/mural cell (clusters 6, 9, 10+15) and fibroblast subpopulations (clusters 1, 2, 8 and 11; summarised Table 1).

## Discussion

Targeting the TME represents an attractive strategy to disrupt tumour-host interactions that promote tumour progression, immune evasion and therapy resistance. While CAF are considered key targets, studies reporting adverse outcomes upon stromal-targeting (Chen et al., 2021; Demircioglu et al., 2020; Özdemir et al., 2014) highlight the functional heterogeneity within the TME and the need for a deeper understanding of fibromuscular cell diversity for effective and safe TME-targeted therapies.

To better characterise the prostatic TME we re-analysed our previously published scRNA-seq PCa dataset identifying multiple fibroblast and SMC/mural cell populations, three of which were associated with favourable (F3/iCAF) or adverse (F5/myCAF and M5/pericytes) clinical outcome. Due to underrepresentation of fibroblasts in this and other publicly available PCa scRNA-seq datasets, we developed an optimised tissue dissociation protocol as a resource for future studies. Compared to two commercial kits, this protocol significantly improved recovery of tissue-resident populations, preserved expression of key fibromuscular cell surface markers and enhanced viable stromal cell isolation revealing differences in stromal composition between benign and malignant tissues and enabling FC-based identification of thirteen distinct stromal clusters. By integrating data from scRNA-seq and FC with immunofluorescence, clusters were annotated as distinct SMC/mural cell and fibroblast subtypes, highlighting the utility of combining orthogonal approaches for profiling heterogeneity of the benign and malignant prostate microenvironment.

Underrepresentation of the fibroblast component in many (PCa) scRNA-seq datasets is an acknowledged limitation (Aparicio et al., 2025) that most likely arises from incomplete release of these ECM-embedded cells, and represented our primary motivation to optimise tissue dissociation as the fundamental step upon which downstream single cell analyses of tissue heterogeneity are based. The dissociation protocol reported here was optimised with respect to the basal digestion buffer, digestion cocktail composition and downstream processing steps. Compared to two commercial formulations and published protocols that employ fewer enzymes and processing steps (Costa et al., 2018; Dominguez et al., 2020; Eich et al., 2018; Quatromoni et al., 2015), the resulting protocol significantly improved viable stromal cell yield and preserved widely employed stromal cell surface markers. Importantly, all pre- and post-digestion steps were performed on ice, with digestion time limited to ≤one hour to minimise dissociation/stress-induced changes (Denisenko et al., 2020; Reichard and Asosingh, 2019). While optimised on benign prostate specimens, the protocol was also effective in dissociating the desmoplastic stroma of PCa tissues. Indeed, FC identified population shifts consistent with tumourigenic hallmarks, including increased endothelial content concordant with tumour neovascularisation, reduced isolation of basal epithelial cells in line with basal cell loss in tumourigenic glands, decreased leukocyte abundance consistent with the immunologically “cold” status of PCa (Brea and Yu, 2025) and enrichment of myCAF/FC cluster 2 in malignant samples. Importantly, our protocol was optimised on small tissue punches, making it suitable not only for cell extraction from primary tumour biopsies but potentially also from soft tissue metastases offering the possibility for clinical exploitation, for example longitudinal tracking for evaluation of therapy response or risk stratification based on stromal subtypes as well as diverse technical and patient-specific primary cell-based applications, such as *ex vivo* drug screening, generation of patient-derived organoids, tissue engineering, progenitor or stem cell therapies, and single-cell multi-omics. Pilot studies using lung cancer specimens similarly improved recovery of tissue-resident cells with intact cell surface marker profiles over existing procedures (data not shown) suggesting our dissociation protocol is potentially of broad applicability.

Due to the absence of unique surface markers for fibroblasts, SMC and mural cells, the FC panel employed combinations of commonly used surface antigens to identify the major prostate cell types and stromal subpopulations recovered. While scRNA-seq of optimally dissociated tissue samples is required for definitive cluster identification, integration with existing scRNA-seq data and immunofluorescent staining enabled annotation of several clusters. Concordant with previous studies (Henry et al., 2018), PDPN and MCAM successfully delineated fibroblast-rich clusters (1, 2, 7, 8) from SMC/mural cells (clusters 6, 11, 10+15), similar to *PDGFRA/MCAM-*based stratification of F and M scRNA-seq clusters. PDPN/MCAM co-expression in FC cluster 9 however constituted a noted exception but was consistent with tissue-based detection of PDPN in PSMC within the fibrotic stroma of benign hyperplastic nodules. Some transcriptionally-defined subpopulations (e.g. proto-myCAF F4 and VSMC/PSMC/pericytes) were less clearly resolved by the surface marker FC panel reflecting their shared lineage and stromal cell plasticity. Discrepancies between transcriptomic and surface marker expression in FC and IF (e.g. *PDGFRA*/CD140a) additionally highlight the challenges in surface marker-based resolution of closely-related stromal cell phenotypes. Future iterations will aim to include additional markers to improve resolution of fibromuscular cell subtypes as well as other cell types e.g. immune cell subsets, to investigate the impact of stromal cell subtypes on immune infiltrates and/or determine how tumour cell genotype influences TME composition. However, the current FC panel represents a strong foundation for stromal characterisation with the potential to evolve into a standalone tool for comprehensive evaluation of cellular heterogeneity of the prostate TME.

Although well established that tumour-associated remodelling of SMC and mural cells exerts clinically-relevant effects (Pederzoli et al., 2023; Taboga et al., 2008), they remain poorly characterised at the molecular level. Within our datasets we identified PSMC and dPSMC as well as two VSMC subtypes and pericytes, whose gene signature correlated with poor prognosis possibly reflecting stabilisation of the tumour vasculature and/or their contribution to the CAF pool via phenotypic switching (Hosaka et al., 2016). FAP^+^ PSMC identified via immunofluorescence in activated benign-adjacent tissues were highly reminiscent of the FAP^+^ subpopulation within FC cluster 6. *FAP* was poorly represented in the scRNA-seq dataset and the current gating strategy did not stratify marker expression levels e.g. decreased MCAM in dPSMC. Thus, it remains to be determined whether FAP^+^ PSMC are analogous to the dedifferentiated, synthetic phenotype of scRNA-seq subcluster M4.

Supported by duplex-ISH, the benign tissue-enriched F1 and F2 scRNA-seq subclusters were identified as benign interstitial and peri-epithelial fibroblasts, respectively. Due to the lack of a unique pan-fibroblast surface marker, these cells could not be unequivocally identified via FC or immunofluorescence. However, given the quiescent nature of benign (non-activated) fibroblasts, as supported by the low number of differentially-expressed genes in subclusters F1/F2, it is plausible that they were represented by one or more of the non-annotated FC clusters (e.g. PDPN^+^MCAM^−^ FC cluster 4+12 and/or non-gated cluster 13+14). Following injury and in fibrotic and cancerous tissues, fibroblasts progress along an activation trajectory comprising an initial pro-inflammatory state that culminates in an ECM-producing myofibroblastic phenotype (Pederzoli et al., 2023; Wietecha et al., 2023). Again, the lack of unique surface markers makes delineating these fibroblast substates a significant challenge. We were thus encouraged by the detection in dissociated tissues and ability of the current FC panel to resolve two of the major CAF substates described to date.

Consistent with previous reports (Hanley et al., 2023), iCAF (scRNAseq cluster F3/FC cluster 8) were prevalent in benign-activated and malignant tissue cores. The F3 gene signature was associated with favourable outcome, potentially reflecting a more immunocompetent TME (Kieffer et al., 2020). Indeed, the FAP⁺CD140b⁺CD90⁺ phenotype of FC cluster 8 aligns with the known role of CD90⁺ fibroblasts in modulating inflammation across benign and malignant contexts (Jenkins et al., 2025; Zeng et al., 2023). Furthermore, high expression of androgen signalling-related genes *PAGE4* and *SRD5A1* in scRNA-seq cluster F3 implies tissue-specific features of prostate iCAF, which as reported recently by us, may further underlie their positive prognostic association as stromal AR suppresses PCa progression (Brunner et al., 2025; Liu et al., 2022).

The myCAF-annotated scRNA-seq cluster F5/FC cluster 2 expressed canonical myCAF markers, in particular FAP and CD105 (ENG), which constituted part of a previously reported prostate CAFÉ (*CTHRC1/ASPN/FAP/ENG*) CAF signature (Wong et al., 2022) with CD105^+^ CAF shown to promote differentiation to the aggressive neuroendocrine state and resistance to AR signalling inhibitors (Kato et al., 2019). Indeed, we recently reported that CD105^+^ primary prostate myCAF express low levels of AR and are proliferatively insensitive to androgen deprivation/AR blockade both *in vitro* and *in vivo* (Brunner et al., 2025). Consistent with multiple reports linking myCAF phenotypes with poor clinical outcome across diverse cancer types (Brunner et al., 2025; Hanley et al., 2023; Kieffer et al., 2020; Li et al., 2021; Mosa et al., 2020; Nicolas et al., 2022; Wong et al., 2022), the F5 gene signature significantly correlated with poor prognostic markers and reduced DFS. Spatially, CD105^+^FAP^+^ cells were enriched in the peri-tumoural space, the established niche of myCAF, where they reportedly contribute to ECM deposition, angiogenesis, and immune exclusion (Brunner et al., 2025; Croizer et al., 2024; Xu et al., 2022).

In summary, the dissociation protocol reported here enables robust recovery of diverse prostate cell types, including fibromuscular subpopulations often under-represented in single-cell studies. Using a multimodal approach, we confirmed distinct fibroblast and mural/SMC phenotypes in the PCa TME, including iCAF and myCAF, underscoring the value of optimised tissue dissociation for capturing stromal heterogeneity. While the stromal response is not currently evaluated in PCa staging, deep learning-based analyses of reactive stromal patterns show promise in improving prognostic accuracy (Ruder et al., 2022). Our findings support such endeavours by defining marker combinations that distinguish PSMC/mural cell types and clinically-relevant fibroblast activation states.

## Declarations

## Ethics approval and consent to participate

Prostate tissue samples were collected from patients undergoing radical prostatectomy at the University Hospital of Innsbruck in accordance with the prior approval from the ethics committee of the Medical University of Innsbruck (1129/2022 and 1072/2018). Use of archived patient-derived FFPE samples was approved by the ethics committee of the Innsbruck Medical University (1129/2022 and 1072/2018). All patients gave written informed consent.

## Consent for publication

All authors have read and consent to this publication.

## Availability of data, materials and code

The source data of the scRNA-seq study re-analysed herein were published elsewhere (Heidegger et al., 2022) and are available in Gene Expression Omnibus (GEO) under accession number GSE193337. Human bulk transcriptomic explant culture dataset is available in ArrayExpress under accession number E-MTAB-13167. Source data are provided in the supplementary file. No new code was developed for this manuscript.

## Authors’ contributions

Conceptualisation: E.D., M.K. and N.S.

Methodology: E.D., E.B., S.S. and N.S.

Software, formal analysis or data curation: E.D., E.B., G.F. and N.S.

Resources: S.S., G.F., Z.T., M.P., I.H. and N.S.

Investigation: E.D., L. Nommensen, L. Neumann and N.S.

Validation: E.D., L. Nommensen, L. Neumann, G.S. and N.S.

Writing – original draft: E.D. and N.S.

Writing – review and editing: S.S., E.B., L. Nommensen, L. Neumann, M.K., G.F., Z.T., M.P., I.H.

Visualization: E.D. and N.S.

Supervision and project administration: N.S.

Funding acquisition: N.S. and M.K.

## Acknowledgements

The authors gratefully acknowledge Gabriele Dobler for assistance with cell culture, Eberhard Steiner for retrieval of clinical data, Sandra Königsdorfer for assistance with tissue collection, Christian Ploner for insightful comments, Andreas Pircher for pre-publication access to the scRNA-seq data, Zoran Culig for infrastructure administration and Iris Magdalena Krainer-Eller, Petra Schumacher, Annabella Pittl, and Corinna Griesbaum for assistance with flow cytometry. The results shown here are in whole or part based upon data generated by the TCGA Research Network (https://www.cancer.gov/tcga). Schematic illustrations were created using BioRender.com.

## Abbreviations

7-AAD: 7-aminoactinomycin D
apCAF: antigen-presenting CAF
AR: androgen receptor
BE: benign
BEC: blood endothelial cells
CA: cancer
CAF: cancer-associated fibroblasts
CI: confidence interval
CK: cytokeratin
DFS: disease-free survival
DMEM: Dulbecco’s modified Eagle medium
DNase: Deoxyribonuclease
DPBS: Dulbecco’s Phosphate-Buffered Saline
dPSMC: dedifferentiated PSMC
ECM: extracellular matrix
EMT: epithelial-mesenchymal transition
EndoC: endothelial cells
EpiC: epithelial cells
FBS: foetal bovine serum
FC: flow cytometry
FFPE: formalin-fixed paraffin-embedded
FMO: fluorescence minus one
FSC-A: forward scatter area
FSC-H: forward scatter height
HBSS: Hank’s Balanced Salt Solution
HE: haematoxylin and eosin
HR: hazard ratio
iCAF: inflammatory CAF
LEC: lymphatic endothelial cells
myCAF: myofibroblastic CAF
P/S: penicillin and streptomycin
panCK: pan-cytokeratin
PCa: prostate cancer
PIN: prostatic intraepithelial neoplasia
PSMC: prostate interstitial smooth muscle cell
scRNA-seq: single cell RNA-sequencing
SMC: smooth muscle cell
TBS: Tris-buffered saline
TME: tumour microenvironment
tSNE: t-distributed stochastic neighbour embedding
VSMC: vascular smooth muscle cell

