## Supplemental Figures for "Optimised dissociation and multimodal profiling of prostate cancer stroma reveal fibromuscular cell heterogeneity with clinical correlates"

### Supplemental Figure legends

**Supplemental Figure 1 Identification of multiple fibromuscular cell populations *in situ***

**(A-B)** Relative distribution of fibroblast (F) and SMC/mural cell (M) subclusters in re-clustered fibromuscular scRNA-seq data (Heidegger et al., 2022) shown per **(A)** patient and **(B)** sample. **(C-D)** Dotplots showing the average expression of the 15 most significant (P.adj) marker genes for each **(C)** fibroblast and **(D)** SMC/mural cell subpopulation. **(E)** Dotplot depicting the expression of published fibroblast and SMC/mural cell gene signatures across fibromuscular subpopulations. Ucell scores were calculated for each gene signature and are represented as a scaled mean per subcluster with dot size inversely proportional to standard deviation (S.D.) to indicate variability within each cluster. **(F)** Dotplot representing average expression of peri-epithelial and interstitial human prostate fibroblast markers (Joseph et al., 2021) across all fibromuscular subpopulations (related to Fig. 1I). **(G)** Violin plot depicting expression levels of selected marker genes demarcating distinct fibroblast (F) and SMC/mural cell (M) subclusters (related to Fig. 1G and 2A). **(H-I)** Heatmaps depicting Ucell scores calculated for published gene signatures and scaled per row for **(H)** fibroblast and **(I)** SMC/mural cell subclusters. Rows are clustered by correlation to published signatures using Ucell scores represented as the mean of each **(H)** fibroblast or **(I)** SMC/mural cell subcluster. **(J-K)** Dual immunohistochemistry for the indicated antibodies whereby font colour corresponds to staining colour and depicting intense CNN1 staining in the outermost VSMC layer of **(J)** arteries and **(K)** veins (black arrowheads). CD105 expression is observed in adventitial fibroblasts, connective tissue surrounding vessels, endothelial cells and myCAF encircling tumour glands (T). Enlarged images of the boxed regions are shown on the right of the parental image. Source data for panels (E) and (H-I) are provided in the Source Data file.

**Supplemental Figure 2 Duplex-ISH reveals distinct prostate fibroblast populations**

**(A-B)** Duplex *in situ* hybridisation of consecutive human benign-adjacent **(A)** and PCa **(B)** tissue sections using the RNA probes indicated, whereby font colour corresponds to staining colour. Sections were counterstained using haematoxylin. Boxed regions are shown enlarged beneath the parental image. Scale bars represent 50 µm (top panels) or 25 µm (middle and lower panels). Images are representative of four independent experiments using tissue sections from three different patients. Black arrowheads denote co-expression of *WNT2* and *C7.* **(C)** Representative images of positive *PPIB* (blue) and *POLR2A* (red) and negative (dapB) control probes run in parallel with experiments shown in (A-B).

**Supplemental Figure 3 Expression of fibromuscular signature genes in TCGA-PRAD cancer and normal prostate tissue cohorts**

**(A-D)** Expression levels (combined z-score) of the most significantly upregulated subpopulation-specific genes in the TCGA-PRAD cancer cohort for the indicated fibroblast (F) and SMC/mural cell (M) subpopulations. Boxplots were used to compare **(A)** Gleason score, **(B)** pathological T stage, **(C)** biochemical recurrence and **(D)** N stage. The R package ggsignif (Ahlmann-Eltze and Patil, 2021) was used to perform T-tests. Statistical significance is denoted NS., not significant; *, $P$<0.05; **, $P$<0.01; ***, $P$<0.001. Source data for panels (A-D) are provided in the Source Data file.

**Supplemental Figure 4 Optimisation of tissue dissociation protocols, comparison of cell diameter and flow cytometry analysis**

**(A)** Viable cell yield/mg tissue under different dissociation conditions (black and grey boxes) during establishment of the optimised dissociation protocol (blue boxes). Bars represent the median ± interquartile range whereby the number of biological replicates (n) is stated beneath each condition. Statistical significance was determined using Brown-Forsythe and Welch ANOVA tests with Dunnett's T3 correction for multiple comparisons. **(B)** Dissociated cell diameter per protocol calculated via ImageJ (as described in Methods) and stratified according to size as indicated. Bars denote median ± interquartile range of 4 biological replicates per protocol. Statistical significance was determined using Brown-Forsythe and Welch ANOVA tests with Dunnett T3 correction for multiple comparisons. **(C)** Flow cytometry histograms for each marker from a representative experiment illustrating overlap of fluorescence signal distribution across all tissue dissociation protocols. Signal intensity is normalised to mode on the Y-axis to facilitate direct comparison of histogram profiles. **(D-I)** Flow cytometry analysis of each tissue dissociation protocol showing the percentage of **(D)** dead, debris, viable and single viable cells in total counts, **(E)** leukocytes, epithelial cells (EpiC), endothelial cells (EndoC) and $CD31^-$ stroma within the single viable cell gate, **(F-G)** EpiC and EndoC further distinguished by PDPN expression into non-basal or basal EpiC, and BEC or LEC as a percentage of **(F)** single viable cells or **(G)** their respective parent gate (EpiC or EndoC), **(H-I)** cells positive for each stromal marker expressed as a percentage of **(H)** single viable cells or **(I)** the CD31- stroma. Bars represent the median ± interquartile range. Statistical significance was determined using (D-G) mixed-effects model with the Geisser-Greenhouse correction and Dunnett correction for multiple comparisons or (H-I) 2-way ANOVA with Dunnett's correction for multiple comparisons. (D-I) Data are derived from multiple independent experiments using tissue samples from different patients whereby n = 4-6 (Miltenyi), 3-6 (BD 60min and 30 min) or 27-29 (Optimised).

**Supplemental Figure 5 Histopathological staining of eight patient cohort**

Dual immunohistochemistry for the basal epithelial cell marker p63 (brown) and tumour cell marker AMACR (red) for histopathological assessment of 4 mm biopsy cores derived from the eight patient-matched cohort. For each patient up to two benign (BE1 or BE2) biopsy cores and one cancerous (CA) biopsy core were sampled. Scale bars represent 500 µm at 20x magnification.

**Supplemental Figure 6 Identification and characterisation of $CD31^-$ stromal clusters via tSNE and flow cytometry gating**

**(A-B)** tSNE dimensionality reduction analysis of $CD31^-$ stroma from flow cytometry data based on the stromal markers MCAM, CD140b, CD90, PDPN, FAP **(A)** without any overlay and **(B)** overlaid with the 16 (unmerged) clusters identified using XShift. **(C-D)** Stacked bar plots depicting the relative distribution of clusters as a percentage of **(C)** the tSNE or **(D)** sample counts according to benign (BE) vs. cancer (CA) histopathology status of the corresponding tissue core. **(E-F)** Relative distribution of **(E)** BE and CA sample counts with overall BE and CA distribution marked as a dotted line and **(F)** patient samples across the 16 clusters. **(G)** Expression of stromal surface markers across the 16 clusters. "Combine" denotes clusters that were subsequently merged into one cluster due to their similar marker expression profiles. **(H)** Relative distribution of the 13 combined clusters across the eight patients. **(I)** Expression of the seven stromal surface markers across the 13 combined clusters. **(J)** Stacked bar plot showing the hierarchical marker distribution within FC-gated cluster 7. The first bar indicates the proportion of $MCAM^+$ and $MCAM^-$ cells within the cluster. Subsequent bars show the distribution of $FAP^+$ and $FAP^-$ populations within each MCAM subset, followed by the relative expression of CD105 within the $FAP^+$ and $FAP^-$ fractions. **(K)** Gating strategy used to identify gated clusters from $CD31^-$ stromal flow cytometry data whereby **(i)** gating for the stromal markers CD140b and PDPN results in **(ii)** a $CD140b^-/PDPN^+$ population, which further subdivided into $CD105^+/FAP^+$ (Cluster 2) and $CD105^-/FAP^-$ (Clusters 4,12), **(iii)** a $CD140b^+/PDPN^+$ population, which includes $CD140a^-/CD90^+$ (Cluster 8), $CD140a^+/CD90^-$

(Cluster 7) and CD140a$^{-}$/CD90$^{-}$ (gated in vi), **(iv)** a CD140b$^{-}$/PDPN$^{-}$ population, which further subdivided into CD140a$^{-}$/CD90$^{+}$ (Cluster 3) and CD140a$^{+}$/CD90$^{-}$ (Cluster 5), **(v)** a CD140b$^{+}$/PDPN$^{-}$ population, which includes MCAM$^{-}$/CD90$^{+}$ (Cluster 11), MCAM$^{+}$/CD90$^{+}$ (Cluster 6), and MCAM$^{+}$/CD90$^{-}$ (Clusters 10,15) and **(vi)** the CD140a$^{-}$/CD90$^{-}$ population from (iii), which further separated into MCAM$^{-}$/FAP$^{+}$ (Cluster 1) and MCAM$^{+}$/FAP$^{-}$ (Cluster 9). The gates were set based on FMO and unstained controls.

**Supplemental Figure 7 Distribution of PDPN immunostaining in the benign and malignant prostate**

**(A-B, D-E)** Immunofluorescent staining of prostate tissue sections of indicated pathology using the antibodies stated whereby font colour denotes pseudocolouring in the merged images. Boxed regions are shown to the right **(A)** or beneath **(B, D)** the parental image. Images are representative of at least three independent experiments using tissues derived from at least five patients. **(A)** Immunostaining of the PSMC and mural cell markers CCDC102B and MCAM reveal strong membrane MCAM immunopositivity of PSMC in benign-adjacent regions (BE, marks benign gland) but a considerable decrease within the tumour core, which still contains CCDC102B$^{+}$ PSMC. **(B)** Immunofluorescent staining of PDPN (red) and CD105 (white) in benign prostate tissue confirming PDPN expression by basal prostate epithelial cells that form a single, continuous layer at the basolateral surface of benign glands. Blood vessels with CD105$^{+}$PDPN$^{-}$ vascular endothelial cells (orange hashtags) are readily distinguished from CD105$^{-}$PDPN$^{+}$ lymphatic endothelial cells (white asterisks). **(C)** Images from the Human Protein Atlas (version 24.0) of immunohistochemical staining of prostate tissues exhibiting low-grade PCa using the anti-PDPN antibody CAB008376 (https://www.proteinatlas.org/ENSG00000162493-PDPN/cancer/prostate+cancer#img). Boxed regions are shown enlarged beneath the parental image. **(D)** Immunofluorescent staining highlighting heterogeneous non-vessel-associated co-expression of MCAM and CD105 (white arrowheads) potentially indicative of pericyte-/SMC-derived myCAF. **(E)** Immunofluorescent staining of a benign-adjacent prostate tissue section exhibiting signs of activation (as indicated by FAP$^{+}$ epithelial cells and PSMC) highlighting vessel-associated co-expression of MCAM, FAP and CD105 in pericytes.

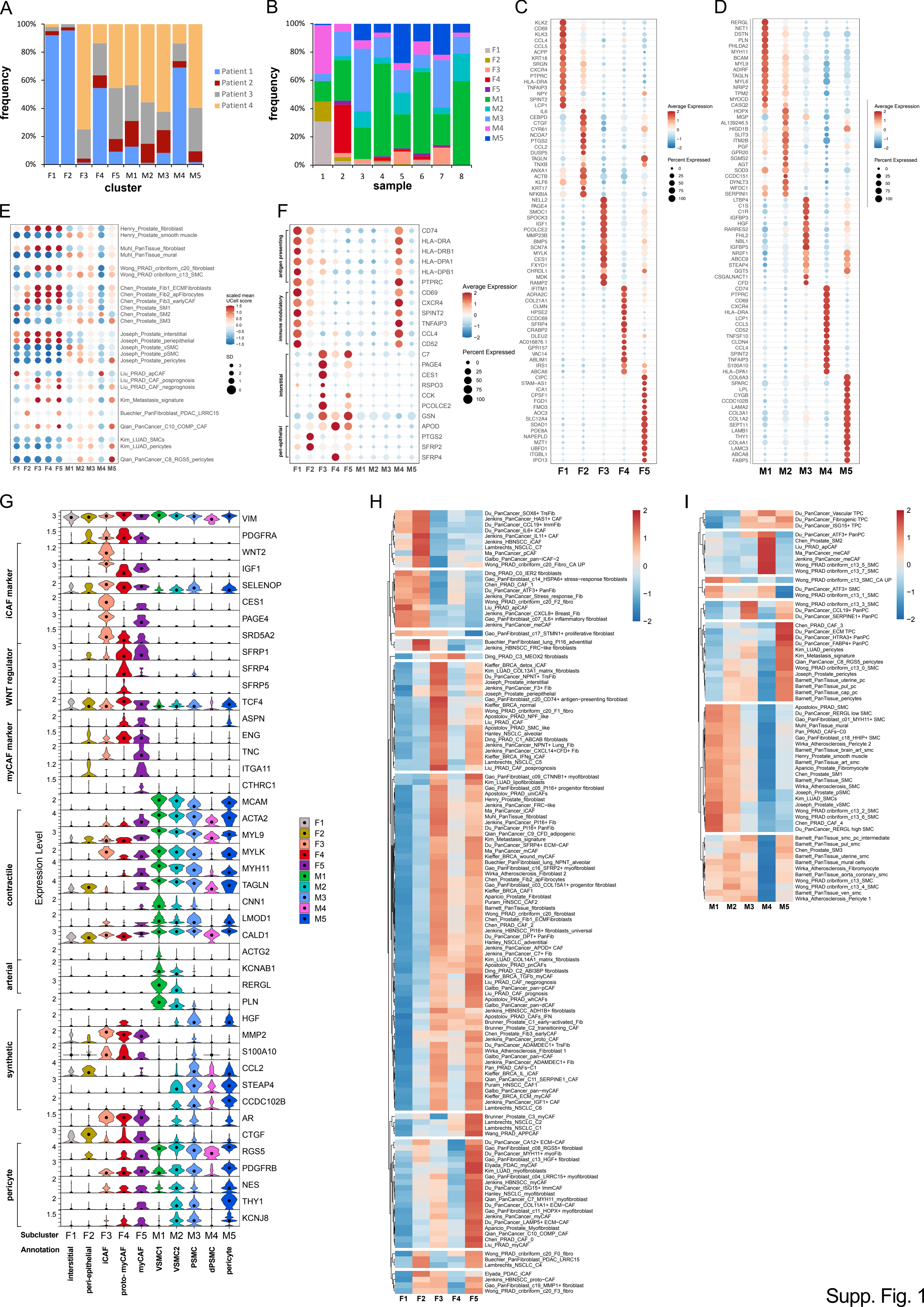
A
frequency
cluster
Patient 1
Patient 2
Patient 3
Patient 4
F1 F2 F3 F4 F5 M1 M2 M3 M4 M5
B
frequency
sample
1 2 3 4 5 6 7 8
F1 F2 F3 F4 F5 M1 M2 M3 M4 M5
C
Average Expression
Percent Expressed
KLK2 CD69 KLK3 CCL4 CCL5 ACPP KRT18 SRGN CXCR4 PTPRC HLA-DRA TNFAIP3 NPY SPINT2 LCP1 IL6 CEBPD CTGF CYR61 NCOA7 PTGS2 CCL2 DUSP5 TAGLN TNXB ANXA1 ACTB KLF6 KRT17 NFKBIA NELL2 PAGE4 SMOC1 SPOCK3 IGF1 PCOLCE2 MMP23B BMP5 SCN7A MYLK CES1 FXYD1 CHRDL1 MDK RAMP2 IFITM1 ADRA2C COL21A1 CLMN HPSE2 CCDC69 SFRP4 CRABP2 DLEU2 AC016876.1 GPR157 VAC14 ABLIM1 IRS1 ABCA6 CIPC STAM-AS1 ICA1 CPSF1 FGD1 FMO3 AOC3 SLC12A4 SDAD1 PDE8A NAPEPLD MZT1 UBFD1 ITGBL1 IPO13
F1 F2 F3 F4 F5
D
RERGL NET1 DSTN PLN PHLDA2 MYH11 BCAM MYL9 ADIRF TAGLN MYL6 NRIP2 TPM2 MYOCD CASQ2 HOPX MGP AL139246.5 HIGD1B SLIT3 ITM2B PGF GPR20 SGMS2 AGT SOD3 CCDC151 DYNLT3 WFDC1 SERPINI1 LTBP4 C1S C1R IGFBP3 HGF RARRES2 FHL2 NBL1 IGFBP5 NR2F1 ABCC9 STEAP4 GGT5 CSGALNACT1 CFD CD74 PTPRC CD69 CXCR4 HLA-DRA LCP1 CCL5 CD52 TNFSF10 CLDN4 CCL4 SPINT2 TNFAIP3 S100A10 HLA-DPA1 COL6A3 SPARC LPL CYGB CCDC102B LAMA2 COL3A1 COL1A2 SEPT11 LAMB1 THY1 COL4A1 LAMC3 ABCA8 FABP5
M1 M2 M3 M4 M5
E
Henry_Prostate_fibroblast
Henry_Prostate_smooth muscle
Muhl_PanTissue_fibroblast
Muhl_PanTissue_mural
Wong_PRAD_cribriform_c20_fibroblast
Wong_PRAD cribriform_c13_SMC
Chen_Prostate_Fib1_ECMFibroblasts
Chen_Prostate_Fib2_apFibrocytes
Chen_Prostate_Fib3_earlyCAF
Chen_Prostate_SM1
Chen_Prostate_SM2
Chen_Prostate_SM3
Joseph_Prostate_interstitial
Joseph_Prostate_periepithelial
Joseph_Prostate_vSMC
Joseph_Prostate_pSMC
Joseph_Prostate_pericytes
Liu_PRAD_apCAF
Liu_PRAD_CAF_posprognosis
Liu_PRAD_CAF_negprognosis
Kim_Metastasis_signature
Buechler_PanFibroblast_PDAC_LRRC15
Qian_PanCancer_C10_COMP_CAF
Kim_LUAD_SMCs
Kim_LUAD_pericytes
Qian_PanCancer_C8_RGS5_pericytes
scaled mean UCell score
SD
F1 F2 F3 F4 F5 M1 M2 M3 M4 M5
F
antigen presenting
immune modulatory
interstitial
peri-epithelial
CD74 HLA-DRA HLA-DRB1 HLA-DPA1 HLA-DPB1 PTPRC CD69 CXCR4 SPINT2 TNFAIP3 CCL4 CD52 C7 PAGE4 CES1 RSPO3 CCK PCOLCE2 GSN APOD PTGS2 SFRP2 SFRP4
Average Expression
Percent Expressed
F1 F2 F3 F4 F5 M1 M2 M3 M4 M5
G
iCAF marker
WNT regulator
myCAF marker
contractile
arterial
synthetic
pericyte
Expression Level
VIM PDGFRA WNT2 IGF1 SELENOP CES1 PAGE4 SRD5A2 SFRP1 SFRP4 SFRP5 TCF4 ASPN ENG TNC ITGA11 CTHRC1 MCAM ACTA2 MYL9 MYLK MYH11 TAGLN CNN1 LMOD1 CALD1 ACTG2 KCNAB1 RERGL PLN HGF MMP2 S100A10 CCL2 STEAP4 CCDC102B AR CTGF RGS5 PDGFRB NES THY1 KCNJ8
Subcluster F1 F2 F3 F4 F5 M1 M2 M3 M4 M5
Annotation interstitial peri-epithelial iCAF proto-myCAF myCAF VSMC1 VSMC2 PSMC dPSMC pericyte
H
F1 F2 F3 F4 F5
I
M1 M2 M3 M4 M5

Supp. Fig. 1

J

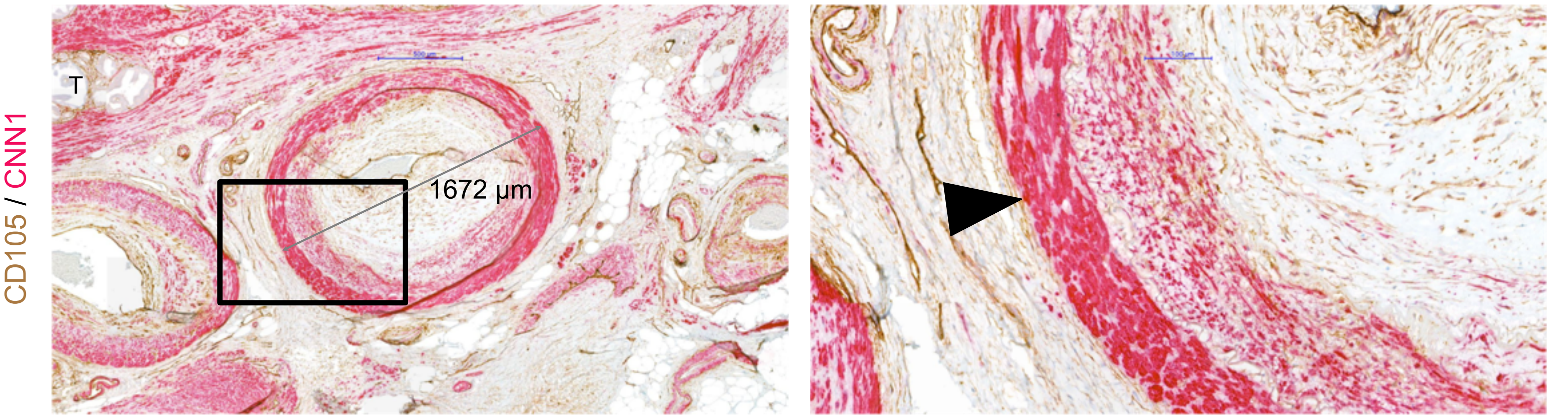


K

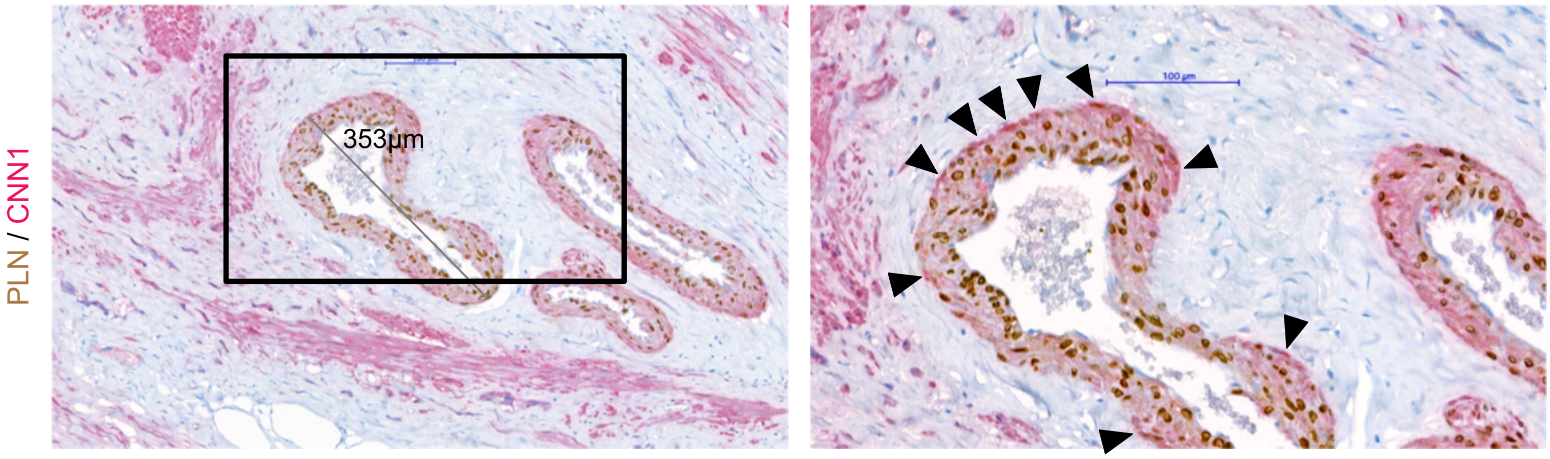


Supp. Fig. 1 cont.

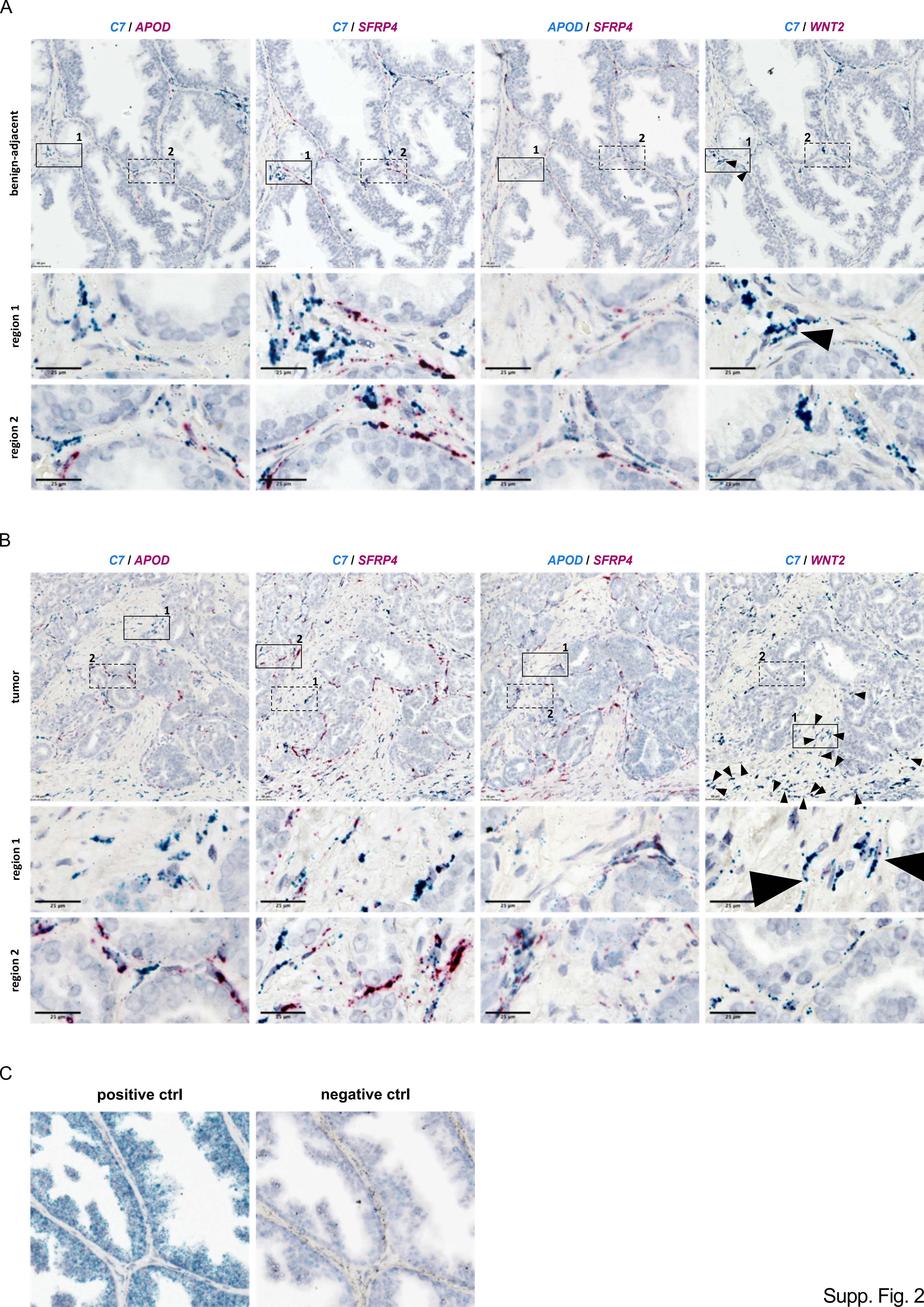
A
C7 / APOD
C7 / SFRP4
APOD / SFRP4
C7 / WNT2
benign-adjacent
region 1
region 2
25 µm
B
C7 / APOD
C7 / SFRP4
APOD / SFRP4
C7 / WNT2
tumor
region 1
region 2
25 µm
C
positive ctrl
negative ctrl

Supp. Fig. 2

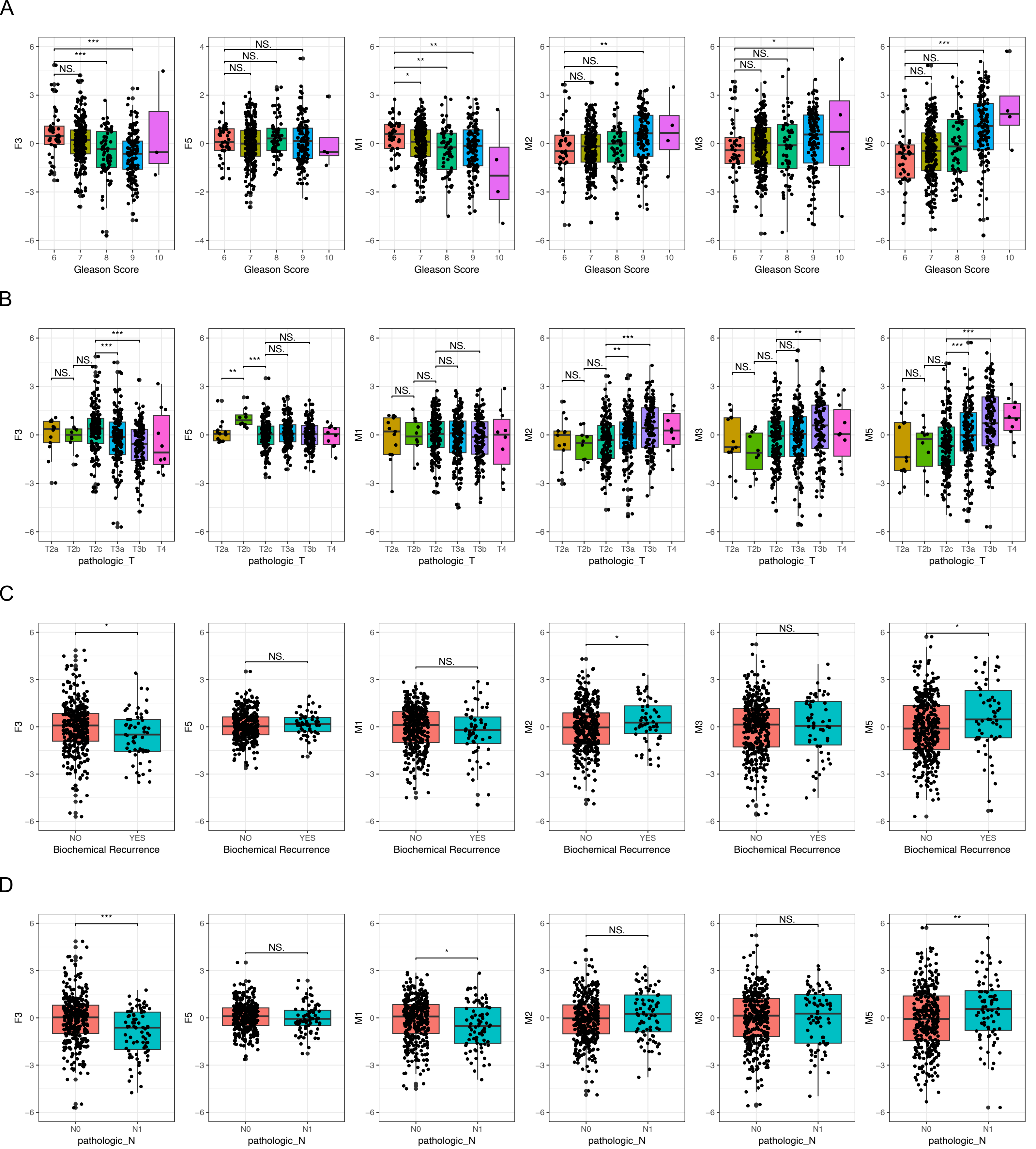
A
F5
F3
M1
M2
M3
M5
Gleason Score
6
7
8
9
10
B
pathologic_T
T2a
T2b
T2c
T3a
T3b
T4
C
Biochemical Recurrence
NO
YES
D
pathologic_N
N0
N1
NS.
*
**
***

Supp. Fig. 3

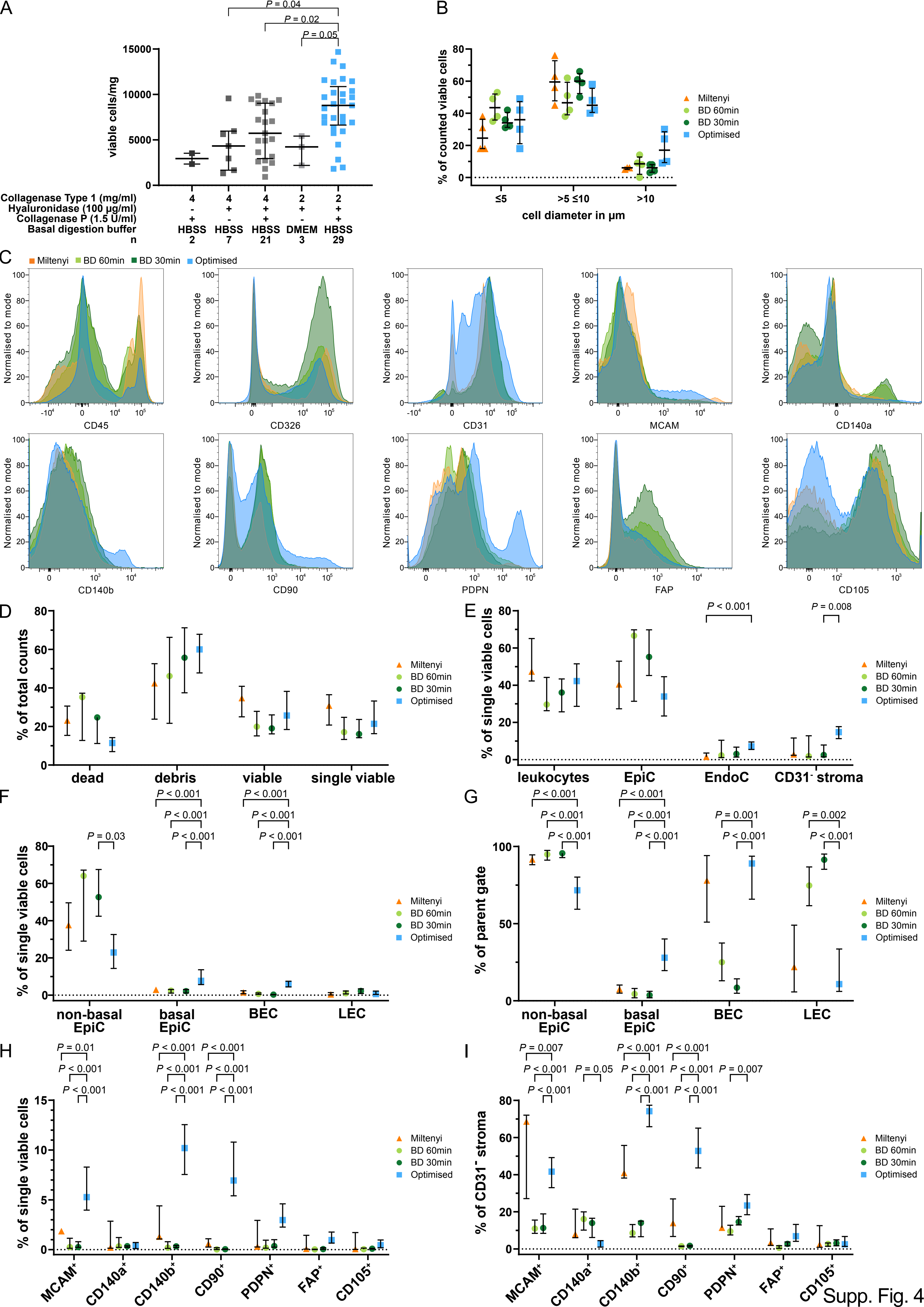
A
viable cells/mg
P = 0.04
P = 0.02
P = 0.05
Collagenase Type 1 (mg/ml)
Hyaluronidase (100 µg/ml)
Collagenase P (1.5 U/ml)
Basal digestion buffer
n
HBSS
DMEM
B
% of counted viable cells
cell diameter in µm
≤5
>5 ≤10
>10
Miltenyi
BD 60min
BD 30min
Optimised
C
Normalised to mode
CD45
CD326
CD31
MCAM
CD140a
CD140b
CD90
PDPN
FAP
CD105
D
% of total counts
dead
debris
viable
single viable
E
% of single viable cells
leukocytes
EpiC
EndoC
CD31⁻ stroma
P < 0.001
P = 0.008
F
non-basal EpiC
basal EpiC
BEC
LEC
P = 0.03
G
% of parent gate
P = 0.001
P = 0.002
H
MCAM⁺
CD140a⁺
CD140b⁺
CD90⁺
PDPN⁺
FAP⁺
CD105⁺
P = 0.01
I
% of CD31⁻ stroma
P = 0.007
P = 0.05
Supp. Fig. 4

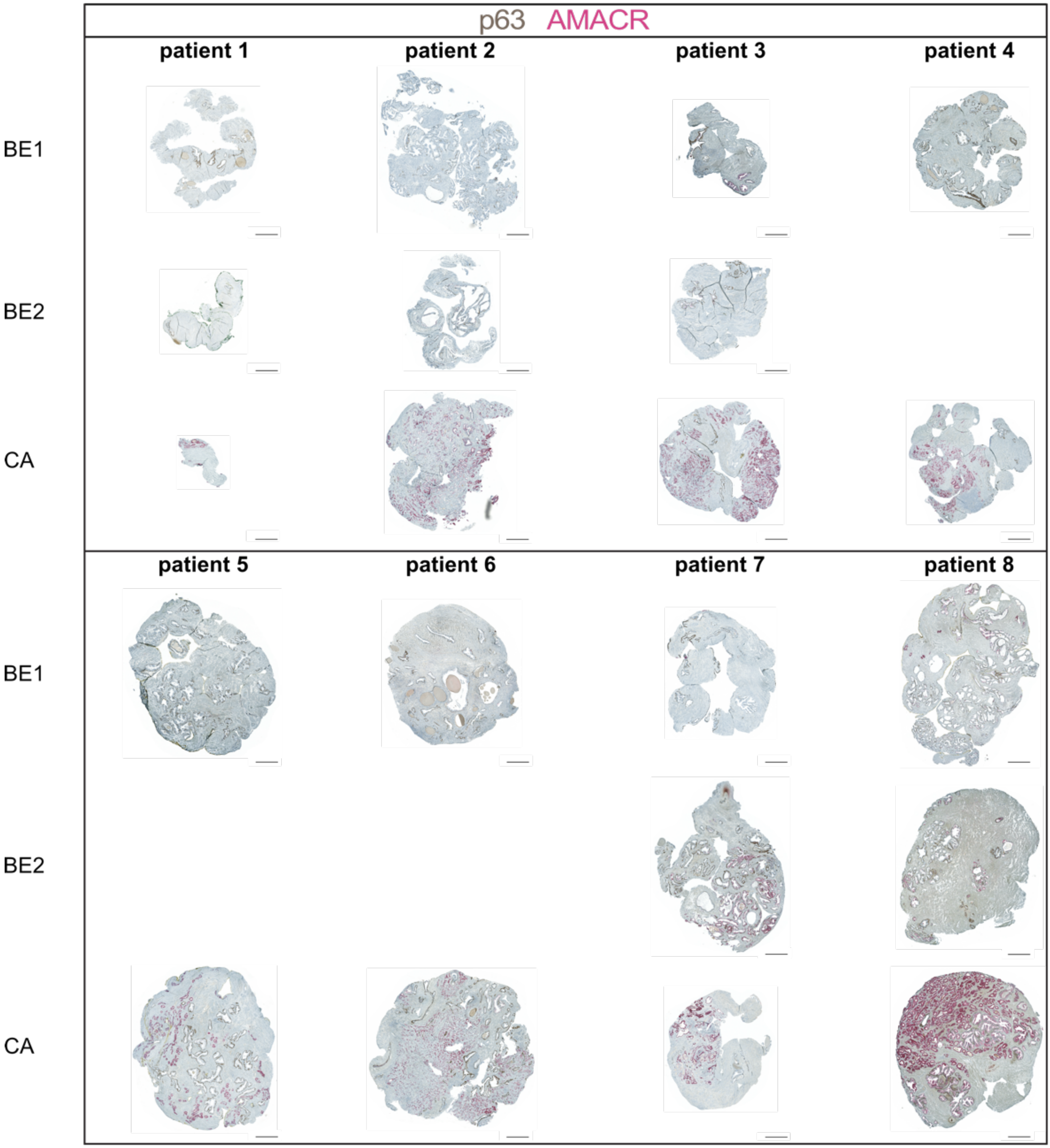
p63 AMACR
patient 1
patient 2
patient 3
patient 4
BE1
BE2
CA
patient 5
patient 6
patient 7
patient 8
BE1
BE2
CA

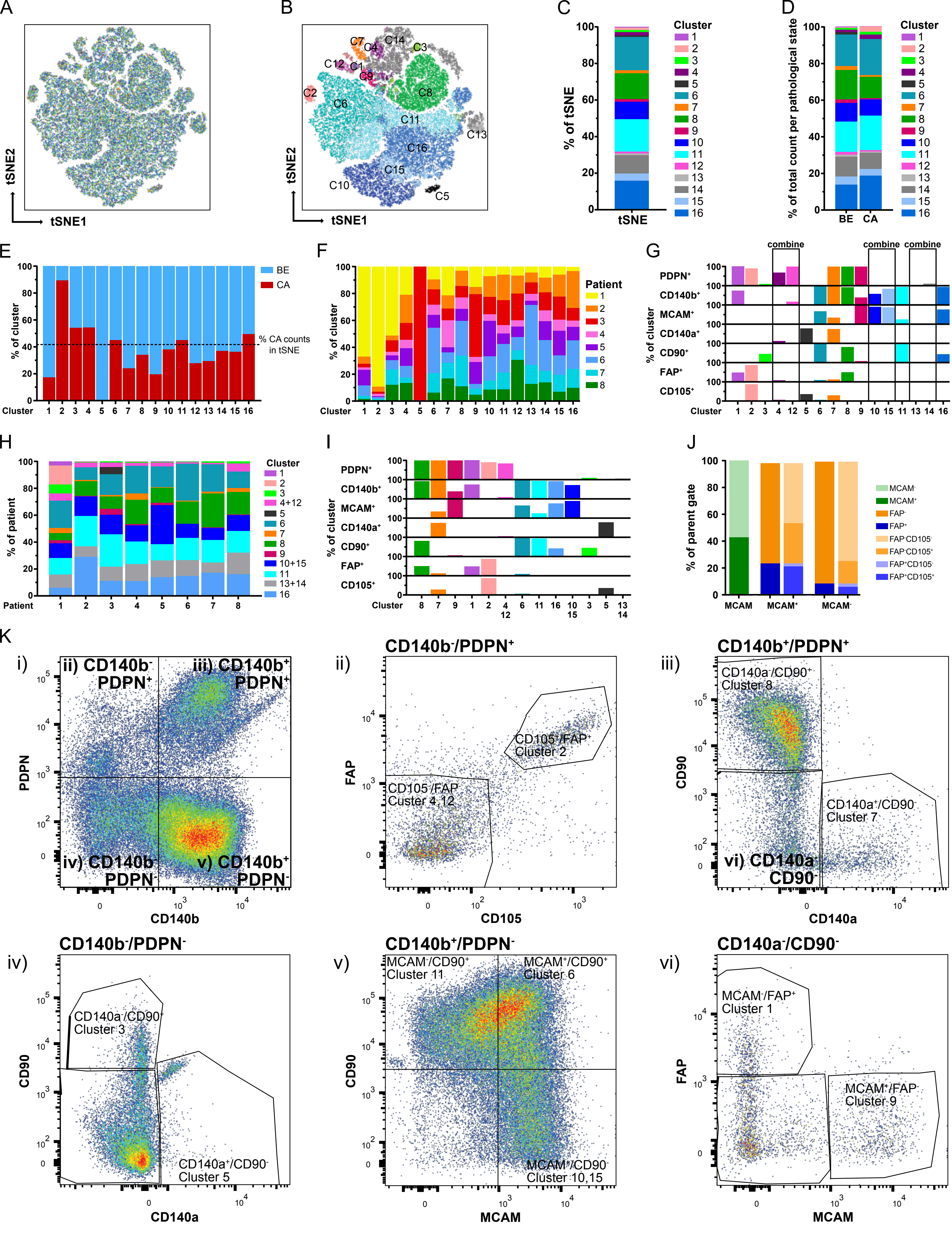
A
tSNE2
tSNE1
B
C7 C4 C14 C3
C12 C1
C9
C2 C6 C8
C11
C13
C16
C15
C10
C5
C
% of tSNE
tSNE
Cluster
1 2 3 4 5 6 7 8 9 10 11 12 13 14 15 16
D
% of total count per pathological state
BE CA
E
% of cluster
BE
CA
% CA counts in tSNE
Cluster
F
Patient
1 2 3 4 5 6 7 8
G
combine
PDPN+
CD140b+
MCAM+
CD140a+
CD90+
FAP+
CD105+
H
% of patient
Patient
4+12
10+15
13+14
I
J
% of parent gate
MCAM
MCAM+
MCAM-
MCAM-
FAP-
FAP+
FAP-CD105-
FAP-CD105+
FAP+CD105-
FAP+CD105+
K
i)
ii) CD140b- PDPN+
iii) CD140b+ PDPN+
iv) CD140b- PDPN-
v) CD140b+ PDPN-
PDPN
CD140b
ii) CD140b-/PDPN+
CD105+/FAP+ Cluster 2
CD105-/FAP- Custer 4,12
FAP
CD105
iii) CD140b+/PDPN+
CD140a-/CD90+ Cluster 8
CD140a+/CD90- Cluster 7
vi) CD140a- CD90-
CD90
CD140a
iv) CD140b-/PDPN-
CD140a-/CD90+ Cluster 3
CD140a+/CD90- Cluster 5
v) CD140b+/PDPN-
MCAM-/CD90+ Cluster 11
MCAM+/CD90+ Cluster 6
MCAM+/CD90- Cluster 10,15
MCAM
vi) CD140a-/CD90-
MCAM-/FAP+ Cluster 1
MCAM+/FAP- Cluster 9

Supp. Fig. 6

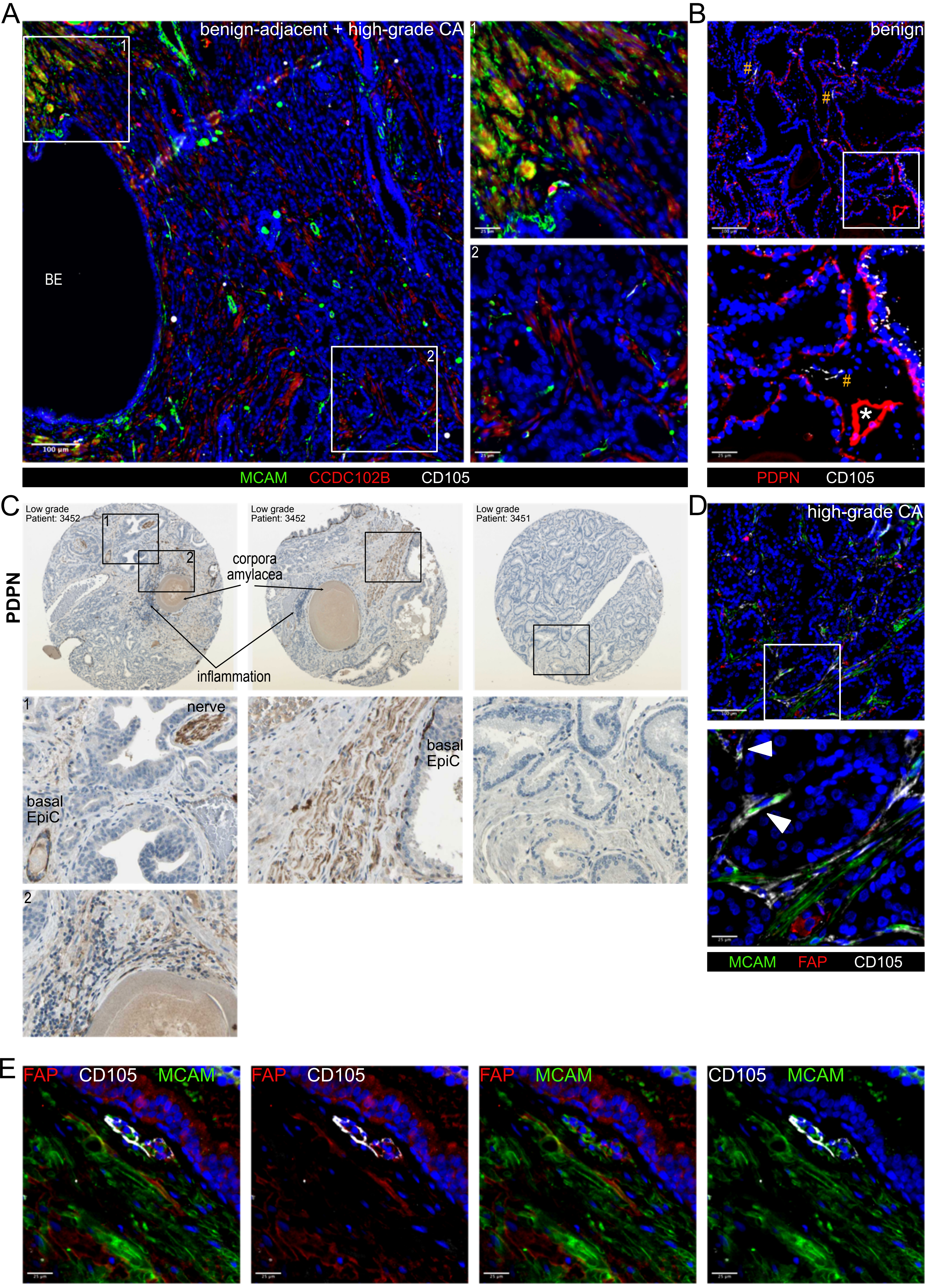
A
benign-adjacent + high-grade CA
BE
100 µm
1
2
25 µm
MCAM CCDC102B CD105
B
benign
#
*
100 µm
PDPN CD105
C
PDPN
Low grade
Patient: 3452
Low grade
Patient: 3452
Low grade
Patient: 3451
corpora
amylacea
inflammation
nerve
basal
EpiC
D
high-grade CA
MCAM FAP CD105
E
FAP CD105 MCAM
FAP CD105
FAP MCAM
CD105 MCAM

Supp. Fig. 7
